# Determinants of human versus mosquito cell entry by the Chikungunya virus envelope proteins

**DOI:** 10.1101/2025.08.25.672233

**Authors:** Xiaohui Ju, William W. Hannon, Tomasz Kaszuba, Caelan E. Radford, Brendan B. Larsen, Samantha S. Nelson, Christopher A. Nelson, Israel Baltazar-Perez, Ofer Zimmerman, Daved H. Fremont, Michael S. Diamond, Jesse D. Bloom

## Abstract

Chikungunya virus (CHIKV) infects both humans and mosquitoes during its transmission cycle. How the virus’s envelope proteins mediate entry in cells from such different species is unclear. MXRA8 is a receptor for CHIKV in mammalian cells, but the receptor(s) in mosquito cells remains unknown. Here we use pseudovirus deep mutational scanning to measure how nearly all amino-acid mutations to the CHIKV envelope proteins affect entry in MXRA8-expressing human and mosquito cells. Most mutations similarly affect entry in both types of cells, and our comprehensive measurements of these effects define functional constraints related to protein folding and fusion activity. However, some mutations differentially affect entry in MXRA8-expressing human cells versus mosquito cells. Sites where mutations specifically impair entry in MXRA8-expressing human cells are often involved in MXRA8 binding, and we hypothesize sites where mutations specifically impair entry in mosquito cells are involved in binding the unknown mosquito receptor(s). We use the deep mutational scanning data to design loss-of-tropism mutant viruses that are impaired in their ability to infect either mosquito cells or MXRA8-expressing human cells. Our findings provide insights into the species-specific determinants of CHIKV cell entry that can help guide receptor identification and vaccine development.

## INTRODUCTION

Chikungunya virus (CHIKV) is an alphavirus that causes musculoskeletal disease and arthritis in humans^1,2^. During its transmission cycle, CHIKV alternates between infecting mammals (humans and non-human primates) and mosquitoes^3,4^. Human outbreaks are increasing due to viral adaptation to additional mosquito species^5–7^ and climate-change driven expansion of mosquito geographic range^8,9^. CHIKV enters both human and mosquito cells using its envelope proteins, which are synthesized as part of a precursor polyprotein that is cleaved into several constituent proteins (capsid, E3, E2, 6K/TF, and E1)^10^. In immature spikes, the envelope proteins consist of heterodimers of p62 (uncleaved E3 and E2) and E1; in mature spikes E3 is cleaved and released from E2, and the resulting E2/E1 heterodimer mediates receptor binding and subsequent fusion of the viral and cell membranes^11,12^. E2 comprises three major domains (A, B, and C), with the A and B domains exposed on the virion surface and contributing to receptor binding and antibody recognition, while the C domain is proximal to the viral membrane^12,13^. The C-terminal region of E2 interacts with the capsid within the virion^14^. E1 is a class II fusion protein composed of three domains (I, II, and III) and contains a fusion loop that inserts into the target-cell membrane during entry^12^. How the envelope proteins enable infection of both human and mosquito cells is unclear. MXRA8 is a receptor for CHIKV in human and mouse cells, although CHIKV can still infect those cells at reduced levels in the absence of MXRA8^15–18^. At the time this study was submitted, receptor in mosquito cells was unknown: an identifiable MXRA8 ortholog is not present in the mosquito genome^15^, and although several candidate mosquito receptors had been proposed, none had been extensively validated^15,18–20^ (see the Discussion for a description of very recent work suggesting lachesin is a mosquito receptor). In addition, several attachment factors (e.g., TIM1, heparan sulfate) can help enable infection at least in cell culture^21–24^.

The envelope proteins are also the antigen targeted by neutralizing antibodies to CHIKV^13,25–27^, and as such are a subject of efforts to develop vaccines^28–30^ and the target of candidate monoclonal antibody prophylactics or therapeutics^25,31,32^. Understanding the functional constraints across these proteins can help inform the design of these countermeasures, which have taken on more importance with the increased global spread of the virus^33,34^. For instance, in the summer of 2025, China experienced its first large CHIKV outbreak, which has reached a scale that has prompted major interventions including efforts to reduce mosquito populations^35,36^.

Here we use pseudovirus deep mutational scanning^37^ to measure how nearly all possible single amino acid mutations to the envelope proteins (E3-E2-6K-E1) affect entry in human and mosquito cells, as well as binding to soluble MXRA8. Our results elucidate the functional constraints on the CHIKV envelope proteins, and define mutations that differentially affect entry in human and mosquito cells.

## RESULTS

### Pseudovirus deep mutational scanning libraries of the CHIKV envelope proteins

To experimentally measure the effects of all amino-acid mutations to the CHIKV envelope proteins (E3-E2-6K-E1), we utilized a recently developed pseudovirus deep mutational scanning system^37,38^. Briefly, this system involves creating libraries of lentiviral particles that each encode a single variant of the envelope proteins in their genome that is also expressed on the surface of the virion, with a unique nucleotide barcode identifying the envelope variant encoded by each virion (**Extended Data Fig. 1a,b**). These pseudoviruses only encode the viral envelope proteins, and so can just undergo a single round of cell entry and are not infectious agents capable of causing disease. They therefore provide a safe way to study the effects of viral protein mutations at Biosafety-level-2.

We created pseudovirus libraries containing nearly all possible single amino-acid mutations to the envelope proteins (E3-E2-6K-E1) from the 181/25 strain of CHIKV (**Extended Data Fig. 1c**), which is an attenuated strain in the Asian lineage^39,40^. Because our libraries are based on a codon-optimized gene for the envelope proteins, they are not expected to produce the TF protein that is generated by ribosomal frameshifting of 6K^41^. We created two independent libraries each designed to contain every possible amino-acid mutation to the 987 residues of the envelope proteins (there are 987 x 19 = 18,753 such mutations). The final libraries contained 134,749 and 124,907 barcoded variants that covered 99.6% and 99.4% of all amino-acid mutations (**Extended Data Fig. 1d**). More than half of the variants contain a single amino-acid mutation, with the others having zero or multiple mutations (**Extended Data Fig. 1e**).

### Effects of mutations on pseudovirus entry into MXRA8-expressing cells

We first measured the effects of all mutations to the envelope proteins on pseudovirus entry into cells expressing the known mammalian receptor for CHIKV, MXRA8^15,16,42^. We created 293T-MXRA8 cells by stably transducing 293T cells (which do not detectably express MXRA8^15^) with a gene encoding human MXRA8 (**Extended Data Fig. 2a**). The titer of pseudovirus expressing the unmutated CHIKV envelope proteins was about 10-fold higher on the 293T-MXRA8 cells than on unmodified 293T cells (**Extended Data Fig. 2b**), consistent with prior data showing that MXRA8 expression enhances CHIKV infection, although some mammalian cells not expressing MXRA8 still can be infected at reduced levels^16–18^. As a control, we confirmed that MXRA8 expression in 293T cells did not affect the titers of pseudovirus expressing the vesicular stomatitis virus G protein (VSV-G) (**Extended Data Fig. 2b**).

To measure the effects of mutations on entry into 293T-MXRA8 cells, we infected these cells with the pseudovirus libraries displaying the CHIKV envelope protein mutants alongside parallel infections of cells with control pseudovirus libraries displaying VSV-G to make their cell-entry independent of CHIKV envelope proteins (**Extended Data Fig. 2c**). We sequenced the barcodes encoded by each pseudovirus variant to quantify the ability of its CHIKV envelope mutant to infect 293T-MXRA8 cells, computing a functional score as the logarithm of the ratio of barcode counts of that variant in the CHIKV envelope-mediated versus VSV-G-mediated infection conditions normalized by the same ratio for the unmutated envelope variants. As expected, envelope variants with only synonymous mutations had wildtype-like functional scores of zero, variants with stop codon mutations had highly negative scores, and variants with amino-acid mutations had scores ranging from wildtype-like to highly negative (**Extended Data Fig. 2d**). To determine the effects of individual mutations on cell entry, we deconvolved the scores for both single- and multi-mutant variants using global epistasis models^43,44^. We performed two technical replicates with each of the two independent libraries, and the measured mutational effects on cell entry were highly correlated both between replicates and libraries (**Extended Data Fig. 2e**). Throughout the rest of this paper, we report the average across all replicates for both libraries.

A full map of how nearly all mutations to the CHIKV envelope proteins affect entry into 293T-MXRA8 cells is shown in **Fig. 1**; the interactive heatmap at https://dms-vep.org/CHIKV-181-25-E-DMS/cell_entry.html can help to more fully explore this large dataset. Mutational effects vary widely across the envelope proteins. Many mutations to E2 and E1 are deleterious for cell entry, consistent with the role of these envelope proteins in the critical processes of receptor binding and cell entry^12,13^ (**Fig. 1**, 2a-b). In contrast, mutations are better tolerated in the E3 protein likely because there is less constraint imposed by its role in chaperoning and stabilizing E2 and E1^45,46^. Most mutations to 6K are well tolerated for pseudovirus cell entry, presumably because its functions as an ion channel and promoter of membrane scission are dispensable in the context of pseudovirus infection in cell culture^47,48^.

**Figure 1.**
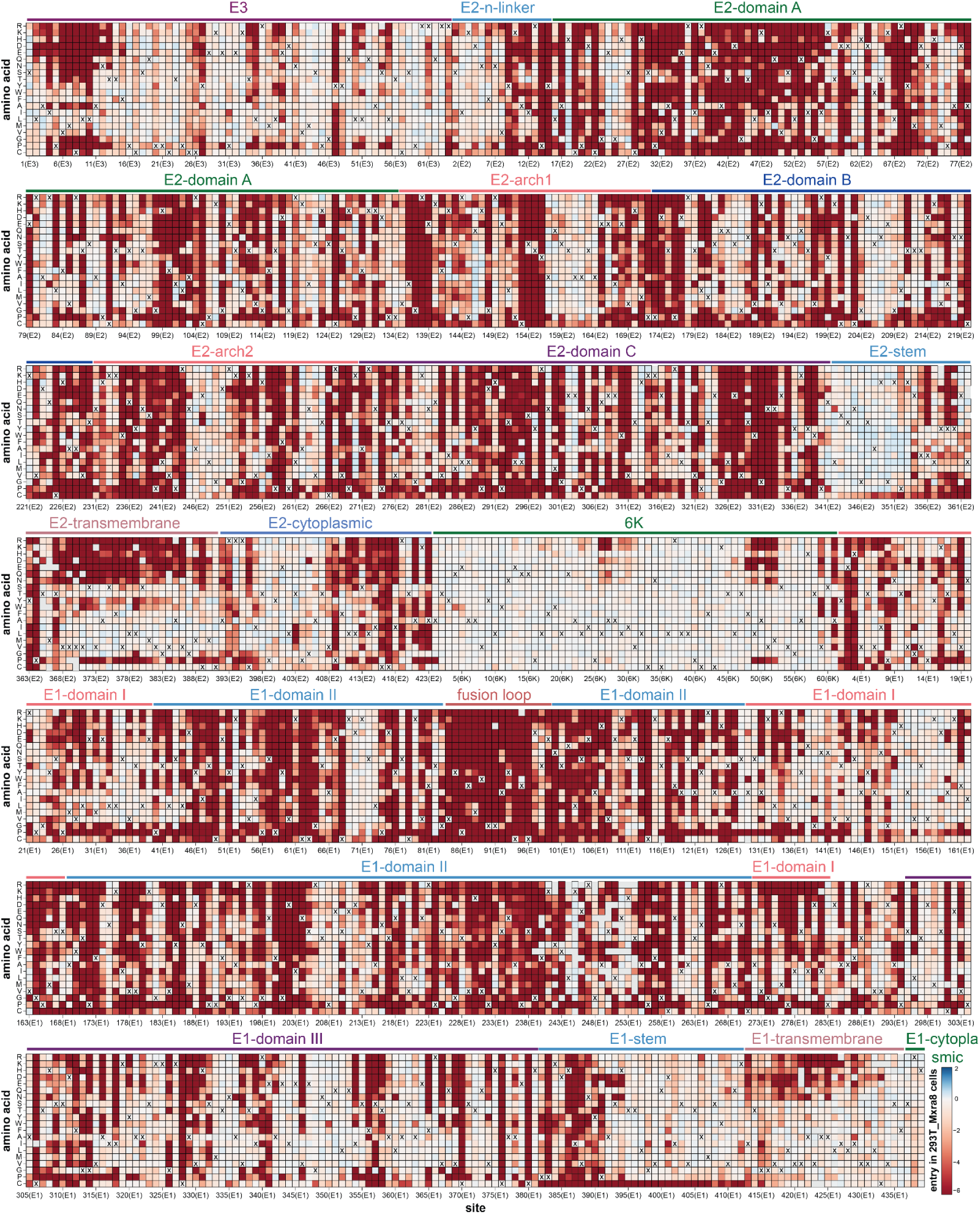
Effects of mutations to the CHIKV envelope proteins on entry in 293T-MXRA8 cells. Each square in the heatmap represents the effect of a mutation, with red indicating impaired entry, white indicating wildtype-like entry, and blue indicating improved entry. The wildtype amino acid in the CHIKV 181/25 strain at each site is indicated with an “X”. The handful of gray squares indicate mutations that were not measured with high confidence in the deep mutational scanning. Protein domains^12^ are labeled above the heatmap. See https://dms-vep.org/CHIKV-181-25-E-DMS/cell_entry.html for an interactive version of this heatmap that is easier to explore than this static version.

**Figure 2.**
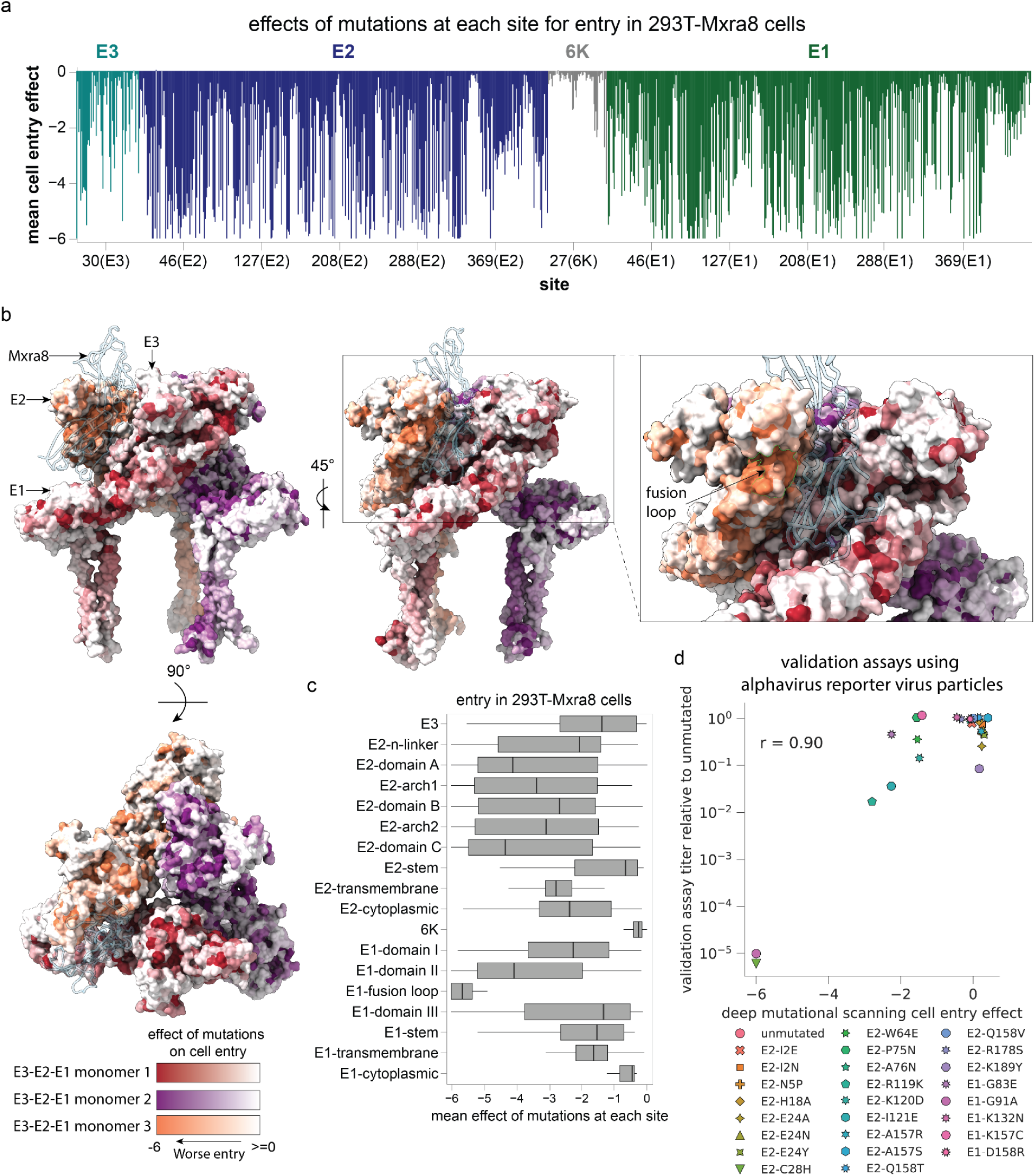
Effects of mutations to different protein domains on entry in 293T-MXRA8 cells. (**a**) Average effect of mutations at each site. (**b**) Trimeric structure of E3-E2-E1 heterotrimer in complex with mouse MXRA8 colored by how mutations affect cell entry (PDB: 6NK7). Each site is colored by the mean effect of mutations at that site, with a different color scale for each monomer of the heterotrimer. Darker colors indicate mutations impair cell entry. MXRA8 is shown in a light blue ribbon. See https://dms-vep.org/CHIKV-181-25-E-DMS/structures.html for interactive visualizations of the mutation effects on the protein structure. (**c**) Distribution of the mean effects of mutations at each site on cell entry for different domains of the envelope proteins. (**d**) Validation assays showing the fold-change in titer in 293T-MXRA8 cells of alphavirus reporter virus particles (RVPs) carrying the indicated mutations versus the mutation effects on cell entry measured by deep mutational scanning. The RVP titers are the mean of three biological replicates for each mutant, and r indicates the Pearson correlation.

Within E2 and E1 there are large differences in the tolerance to different mutations with respect to entry in 293T-MXRA8 cells (**Fig 1**). Some of these patterns in mutational effects can be understood in terms of protein structure and function. For instance, domain C of E2 is highly intolerant to mutations (**Fig. 1**, 2c), probably due to its importance in spike trimerization^12,49^. In contrast, the E2 stem tolerates most mutations (**Fig. 1, 2c**), probably because this region simply needs to connect the E2 ectodomain to the transmembrane domain that anchors spike in the viral membrane^12,50^. The most constrained region of the envelope proteins is the fusion loop of E1 where nearly all mutations are deleterious (**Fig. 1, 2b-c**), consistent with its highly conserved role in mediating fusion of viral and cell membranes^51,52^. Note that in some cases differences between pseudovirus and actual virus could affect the results: for instance, the cytoplasmic region of E2 is relatively tolerant to mutations in our pseudovirus deep mutational scanning but more constrained in natural sequences (**Extended Data Fig. 2f**), probably because this domain normally interacts with the CHIKV capsid protein^14^ that is absent in our pseudovirus system.

To validate the pseudovirus deep mutation scanning results in actual alphavirus virion particles, we used a single-cycle reporter virus particle (RVP) system^53,54^ in which a truncated alphavirus genomic RNA that encodes the Venezuelan equine encephalitis virus (VEEV) attenuated strain TC-83 nonstructural proteins (but not the structural proteins) is packaged into virions bearing capsid and envelope proteins of CHIKV by co-expressing these structural proteins from separate plasmids (**Extended Data Fig. 2g**). These pseudoinfectious virions cannot undergo multi-cycle infection as they do not encode the structural proteins. We constructed RVPs carrying 25 different mutations to the CHIKV envelope proteins that had a range of effects in the pseudovirus deep mutational scanning. The individual RVP titers were correlated highly with the deep mutational scanning measurements (r = 0.90; **Fig. 2d**), demonstrating a correspondence between the effects of mutations to the envelope proteins on entry of lentivirus-based pseudovirus and bona fide alphavirus particles in 293T-MXRA8 cells.

### Effects of mutations on MXRA8 binding

Mutations to the CHIKV envelope proteins could affect entry into MXRA8-expressing cells by altering a variety of protein properties, including MXRA8 binding, membrane fusion activity, and protein folding and stability. To isolate the effects of mutations on MXRA8 binding, we utilized the fact that inhibition of virion cell entry by soluble decoy receptors is proportional to receptor-binding affinity^38,55–57^. First, we confirmed that both soluble human and mouse MXRA8-Fc fusion proteins inhibited infection of 293T-MXRA8 cells by both pseudovirus expressing the CHIKV envelope proteins and authentic CHIKV of the 181/25 vaccine strain (**Fig. 3a**). Mouse MXRA8 was more potent than human MXRA8 at inhibiting infection, consistent with prior work showing stronger binding of the CHIKV envelope proteins to mouse versus human MXRA8^15,51,52^; authentic CHIKV was inhibited by somewhat lower concentrations of MXRA8-Fc than pseudovirus, possibly because authentic virus has a higher density of envelope proteins thereby enabling better MXRA8 binding. We measured the effects of envelope protein mutations on MXRA8 binding by incubating the pseudovirus libraries with different concentrations of soluble mouse or human MXRA8-Fc, also including a VSV-G pseudotyped virus as a standard to convert sequencing counts to the fraction of infectivity retained^37^ (**Extended Data Fig. 3a**). Note that this approach can only measure the effect on MXRA8 binding of envelope protein mutants that retain at least minimal level of cell entry in MXRA8-expressing cells.

**Figure 3.**
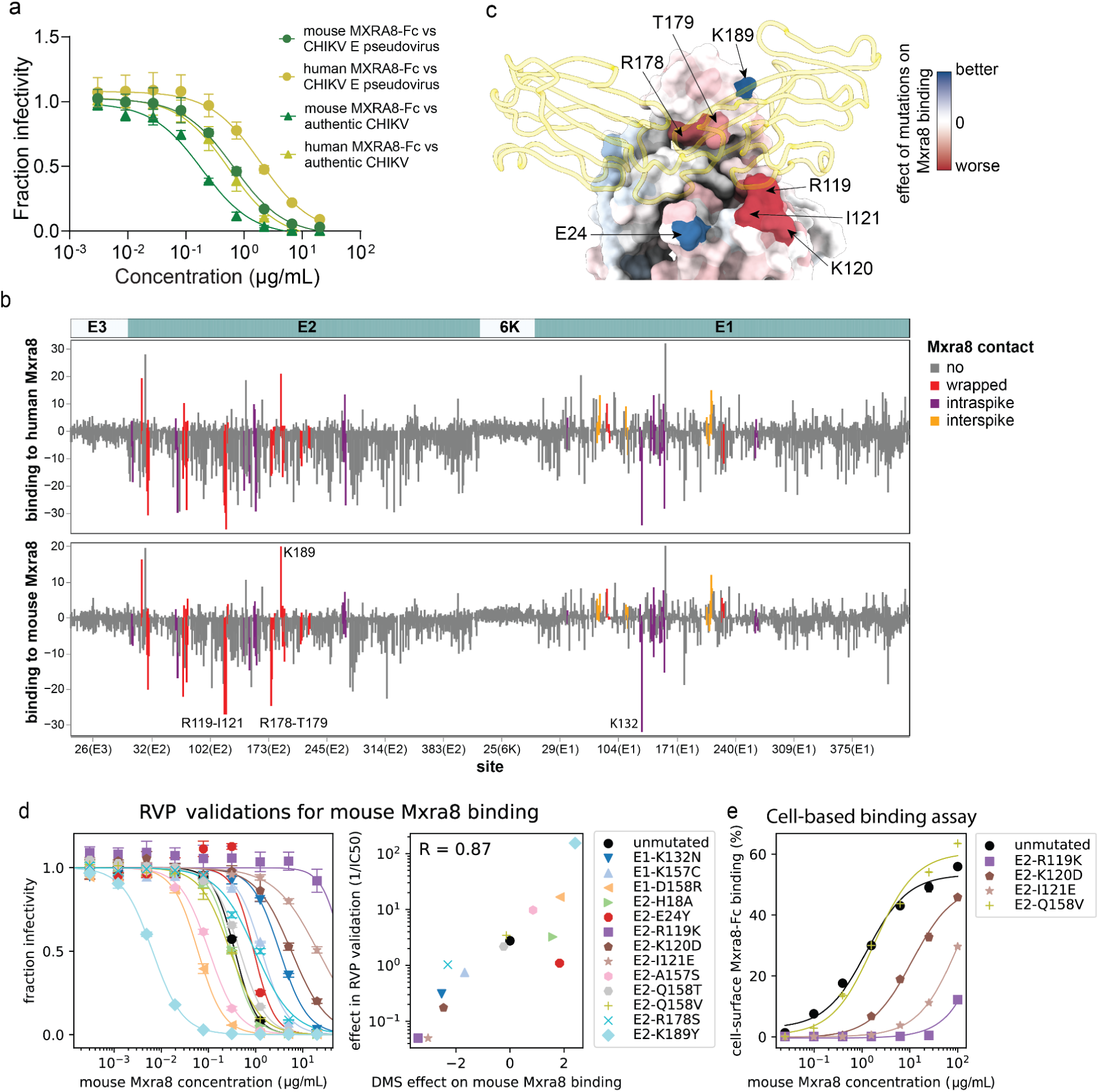
Effects of mutations on binding to MXRA8. (**a**) Neutralization of pseudovirus expressing unmutated CHIKV envelope proteins and authentic CHIKV of the 181/25 vaccine strain by soluble mouse or human MXRA8-Fc. (**b**) Deep mutational scanning measurements of effects of mutations at each site on binding to human (top) or mouse (bottom) MXRA8. At each site, the plot shows the sum of all mutations that increase binding (positive values) and the sum of all mutations that decrease binding (negative values). Sites are colored by whether and how they contact MXRA8 as annotated by Basore et al^51^. Sites not annotated as contact sites^51^ that have an appreciable effect on MXRA8 binding tend to be close to MXRA8 in cryo-EM structures as shown in **Extended Data Fig. 3c**. See https://dms-vep.org/CHIKV-181-25-E-DMS/MXRA8_binding.html for an interactive version of this plot. (**c**) A single E3-E2-E1 monomer from cryo-EM structure of the full trimer (PDB: 6NK7) colored by the site-summed effects of mutations on binding to mouse MXRA8, with mouse MXRA8 shown in the wrapped contact mode with a light gold ribbon. Sites lacking binding measurements (because mutations are highly deleterious for cell entry) are colored light grey. See https://dms-vep.org/CHIKV-181-25-E-DMS/structures.html for interactive visualizations of the mutation effects on the structure. (**d**) Validation assays showing the neutralization of alphavirus RVPs carrying the indicated E1 or E2 mutations by soluble mouse MXRA8-Fc (left panel), and the correlation of the inverse IC50 for the RVP neutralization with the effect of that mutation measured in the deep mutational scanning (DMS) (right panel). The plot shows the mean of three biological replicates for each RVP mutant, and R indicates the Pearson correlation. **(e)** Binding of soluble mouse MXRA8-Fc to 293T cells transfected with unmutated or mutant CHIKV envelope proteins as quantified by flow cytometry. The y-axis shows the percent of the envelope-protein expressing cells that are stained positive for MXRA8 binding at each MXRA8 concentration. See **Extended Data Fig. 3f** for representative flow cytometry plots.

The sites where mutations most strongly affect MXRA8 binding are located primarily in E2 and to a lesser extent E1 (**Fig. 3b**, **Extended Data Fig 4**, and interactive plots at https://dms-vep.org/CHIKV-181-25-E-DMS/MXRA8_binding.html), consistent with structural data showing that E2 is the primary receptor-binding protein with E1 also contributing some MXRA8 contact residues^51,52^. Mutations at most sites in the envelope proteins had similar effects on binding to mouse and human MXRA8 (**Fig. 3b**, **Extended Data Fig. 3b**), consistent with cryo-EM structures showing MXRA8 proteins from these two different species have a similar binding interface with CHIKV envelope proteins^51,52^. We were able to make higher-quality measurements (as assessed by correlations between deep mutational scanning library replicates; see **Methods**) for mouse MXRA8 binding than for human MXRA8 binding, probably due to its greater potency at neutralizing pseudovirus (**Fig. 3a**). Prior structural work has shown that mouse MXRA8 interacts with the E2-E1 heterodimer via three different dimer binding modes termed wrapped, intraspike, and interspike^51^. In our deep mutational scanning, mutations at sites that contact MXRA8 in the wrapped conformation have the largest effects on binding, with mutations at some intraspike contact sites also having appreciable effects and mutations at interspike contact sites having smaller effects (**Fig. 3b**, **Extended Data Fig. 3c and 4**). Mutations at sites that contact MXRA8 in the wrapped binding mode clearly have the largest impact on entry into cells expressing MXRA8 (**Extended Data Fig. 3d**), consistent with structural evidence that this mode makes the largest contribution to MXRA8 binding^51^. The effects of mutations on MXRA8 binding and entry in 293T-MXRA8 cells are only weakly correlated (**Extended Data Fig. 3e**), suggesting that most mutations that impair pseudovirus entry in the 293T-MXRA8 cells (which express high levels of receptor) do so by disrupting folding or fusion activity of the envelope proteins rather than by impairing receptor binding. However mutations in E2 that strongly impair MXRA8 binding are deleterious for entry in 293T-MXRA8 cells (e.g., G118K, R119A, R119S, I121G and I121K; **Extended Data Fig. 3e**).

The sites where mutations most strongly decrease MXRA8 binding include the wrapped E2 contact sites R119-I121 and R178-T179, as well as the E1 intraspike contact site K132 (**Fig. 3b-c**, **Extended Data Fig. 4**). This finding is consistent with the structure of E2-E1 bound to MXRA8, E2 sites R119-I121 interact with human MXRA8 D90-R98 and mutations of Y92, E96 or R98 in human MXRA8 decrease its binding to envelope proteins^52^. Mutations at some of the homologous sites in mouse MXRA8 (D89R, G94R, and R97E) also markedly reduce CHIKV infection in cell lines^51^. There are also some sites where mutations to the envelope proteins increase MXRA8 binding, including E2 K189, which contacts mouse MXRA8 Q83 in the wrapped binding mode (**Fig. 3b-c**, **Extended Data Fig. 4**).

To validate the pseudovirus deep mutational scanning measurements of how mutations affect MXRA8 binding, we used the alphavirus RVP system to measure how 13 different mutations with a range of effects in the deep mutational scanning affected neutralization by soluble mouse MXRA8-Fc in 293T-MXRA8 cells. The effects of the mutations on MXRA8 neutralization of the RVPs was correlated highly with the deep mutational scanning results (**Fig. 3d**). To directly confirm that neutralization of the pseudoviruses by soluble MXRA8-Fc corresponds to its binding to the envelope proteins, we used flow cytometry to demonstrate that three mutations that decreased soluble MXRA8 neutralization (R119K, K120D, I121E) also reduced binding of MXRA8 to cell-surface expressed envelope proteins, while a mutation that had no effect on soluble MXRA8-Fc neutralization (Q158V) also did not impact binding (**Fig. 3e, Extended Data Fig. 3f**).

### Effects of mutations on entry in mosquito cells or human cells expressing the TIM1 attachment factor

To determine how mutations to the envelope proteins affect entry in cells that do not express MXRA8, we examined pseudovirus entry in two additional cell lines. The first of these cell lines was C6/36 cells, which are derived from *Aedes albopictus* mosquitoes^58^ and are widely used in CHIKV studies^59–63^. We also used a previously described^64^ 293T cell line that lacks MXRA8 expression but does express TIM1, which promotes infection of several enveloped viruses including CHIKV by binding to virion-associated phosphatidylserine^18,21,65,66^. However, it is unclear if TIM1-mediated CHIKV infection is relevant in authentic human infections or is just a cell-line phenomenon^67,68^. Expression of TIM1 in 293T cells increased the titers of pseudovirus expressing the unmutated CHIKV envelope proteins by >10-fold, an even greater increase than that caused by MXRA8 expression (**Fig. 4a**). Expression of TIM1 in 293T cells also increased the titers of alphavirus RVPs and replication of CHIKV 181/25, although for these the increase was less than that caused by MXRA8 expression (**Fig. 4a**). Note that 293T cells but not C6/36 cells express heparan sulfate, which also can act as an attachment factor for CHIKV^22–24^ (**Extended Data Fig. 5a**).

**Figure 4.**
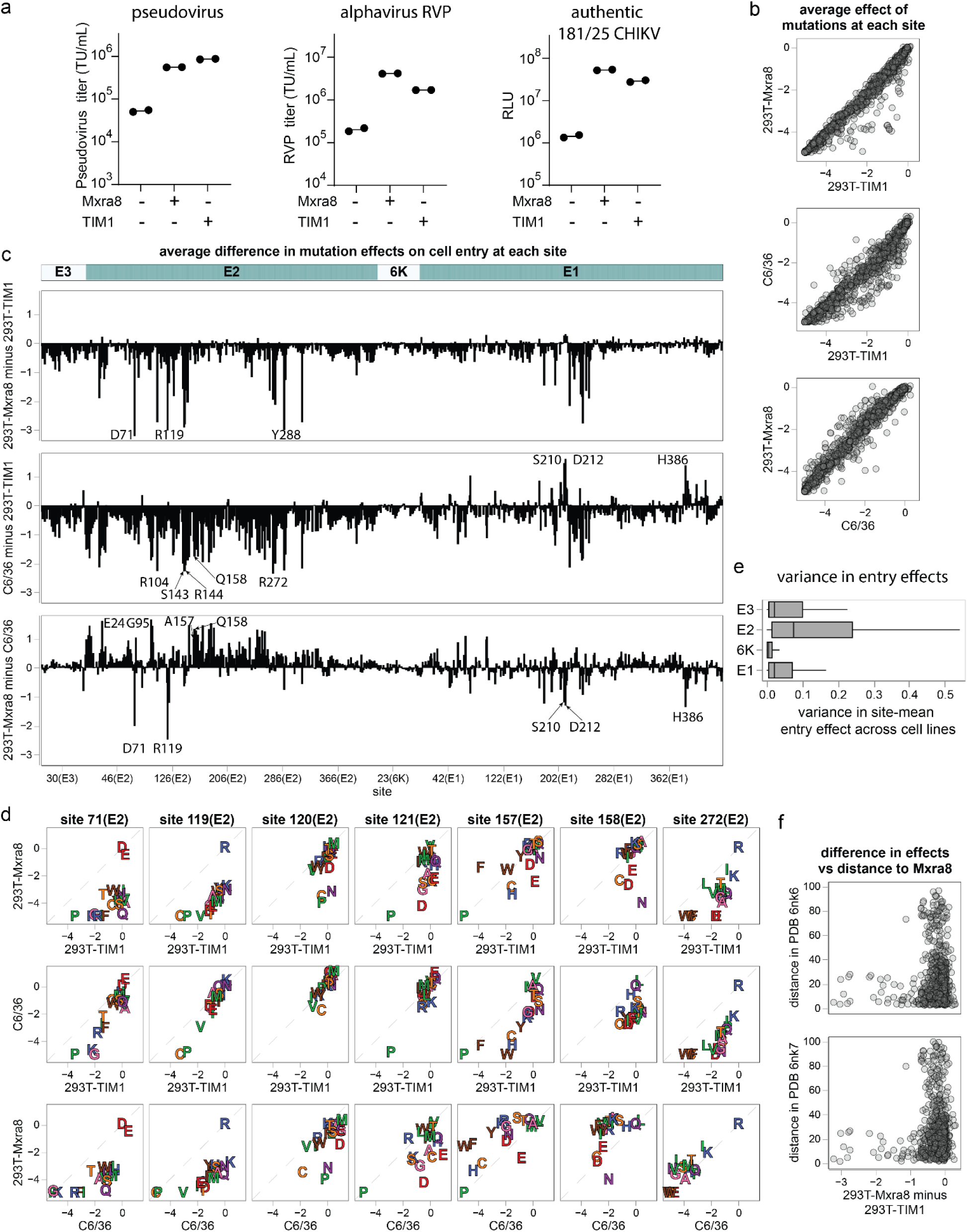
Comparison of effects of mutations on entry in cells expressing MXRA8, mosquito cells, and cells expressing TIM1. (**a**) Titers of pseudoviruses, alphavirus RVPs, and CHIKV 181/25 on 293T, 293T-MXRA8 or 293T-TIM1 target cells. Pseudovirus and RVP titers were quantified as transducing units (TU) per ml; CHIKV 181/25 titers were quantified by the relative light units (RLU) produced by a luciferase reporter in the viral genome at 24 hours after infection. (**b**) Mean effects of mutations at each site on pseudovirus entry in 293T-MXRA8, 293T-TIM1 or C6/36 cells as measured by deep mutational scanning. Each point represents a single protein site. Mutations that have similar effects in each pair of cells are on the diagonal; mutations with different effects are off the diagonal. See https://dms-vep.org/CHIKV-181-25-E-DMS/cell_entry_diffs.html for interactive visualizations of the differences in mutation effects on entry into different cells. (**c**) Differences in effects of mutations at each site on entry in different cells. For each site, the plot shows the summed differences in effects of all mutations in the first cell minus the second cell. For instance, the negative value for site D71 in the top plot indicates mutations at this site are more deleterious for entry in 293T-MXRA8 cells than 293T-TIM1 cells. (**d**) Effects of individual mutations on entry in each cell for some sites with large cell-to-cell differences. In each scatter plot, each letter represents the effect of that amino acid on entry in each of the two indicated cells. Letters are colored by the biophysical properties of the amino acids. **(e)** Variance in effects of mutations at each site on cell entry across the three cell lines, with larger values indicating mutations have larger differences in their effects in different cells. **(f)** Difference in effects of mutations on entry in 293T-MXRA8 minus 293T-TIM1 cells at each site plotted against the distance of the site from MXRA8 in cryo-EM structures (PDB identifiers 6nk6 or 6nk7)^51^. Sites where mutations are appreciably more deleterious for entry in 293T-MXRA8 than 293T-TIM1 cells (negative differences) tend to be close to MXRA8 in the structures.

We used the same deep mutational scanning workflow described above (**Extended Data Fig. 2c**) to measure the effects of mutations on entry in 293T-TIM1 and C6/36 cells. Note that while it is unclear if the pseudoviruses fully integrate their lentiviral genomes in the C6/36 cells, they did robustly produce the non-integrated lentiviral genomes needed for our sequencing-based readout. For both 293T-TIM1 and C6/36 cells, envelope variants with synonymous mutations had wildtype-like functional scores, variants with stop codon mutations had highly negative scores, and variants with amino-acid mutations had scores ranging from wildtype-like to highly negative (**Extended Data Fig. 5b**), a pattern resembling that seen for 293T-MXRA8 cells (**Extended Data Fig. 2d**). The measurements of effects of mutations on entry in each cell line were highly correlated both between technical replicates and independent libraries (**Extended Data Fig. 5c-d**). The full measured effects of all mutations in these two cell lines are shown in **Extended Data Figs. 6 and 7** and the interactive plots at https://dms-vep.org/CHIKV-181-25-E-DMS/cell_entry.html. We validated the pseudovirus deep mutational scanning measurements for these two cell lines for several mutations using alphavirus RVPs (**Extended Data Fig. 5e**).

Most mutations had similar effects on entry in all three cell lines, but at a few sites mutations had distinct effects in the different cells (**Fig. 4b-d** and the interactive comparison at https://dms-vep.org/CHIKV-181-25-E-DMS/cell_entry_diffs.html). In most cases, the pattern was for mutations that are deleterious for entry in either 293T-MXRA8 or C6/36 cells to be tolerated in 293T-TIM1 cells (**Fig. 4b-d**). For instance, many mutations at E2 sites 71, 119-121, and 288 are deleterious for entry in 293T-MXRA8 cells, but often are well tolerated for entry in 293T-TIM1 cells (**Fig. 4b-d)**. Likewise, nearly all mutations at E2 sites 104, 143, 144, 158, and 272 are deleterious for entry in C6/36 cells, but many have little impact on entry in 293T-TIM1 cells (**Fig. 4b-d**). This pattern makes sense in terms of the role of the envelope proteins for entry in each cell line. For all three cells, entry depends on proper folding and membrane fusion by the envelope proteins, and mutations affecting these properties have similar effects in all three cells (**Fig. 4b**). However, the mechanism of virion attachment differs across the cells: attachment to 293T-MXRA8 cells is mediated largely by binding to MXRA8, attachment to C6/36 cells is likely mediated by binding to some unknown receptor, and attachment to 293T-TIM1 cells is mediated by binding of TIM1 to virion-associated phosphatidylserine that is independent of the envelope proteins^65,66^. Indeed, among the four envelope proteins, E2—which is the main determinant of receptor binding—exhibits the greatest variance in the effects of mutations across cell lines (**Fig. 4e**).

The mutations that are more deleterious for entry in 293T-MXRA8 cells than 293T-TIM1 cells are mostly at sites in or near the MXRA8 binding interfaces (**Fig. 4f**, **5**), consistent with the fact that entry into 293T-MXRA8 but not 293T-TIM1 cells involves MXRA8 binding. For example, E2 sites 71, 119, 120 and 121 contact MXRA8 in the wrapped binding mode and E2 site 144 contacts MXRA8 in the intraspike binding mode—and mutations at all these sites are more deleterious for entry in 293T-MXRA8 than 293T-TIM1 cells (**Fig. 4b-d, 5**).

**Figure 5.**
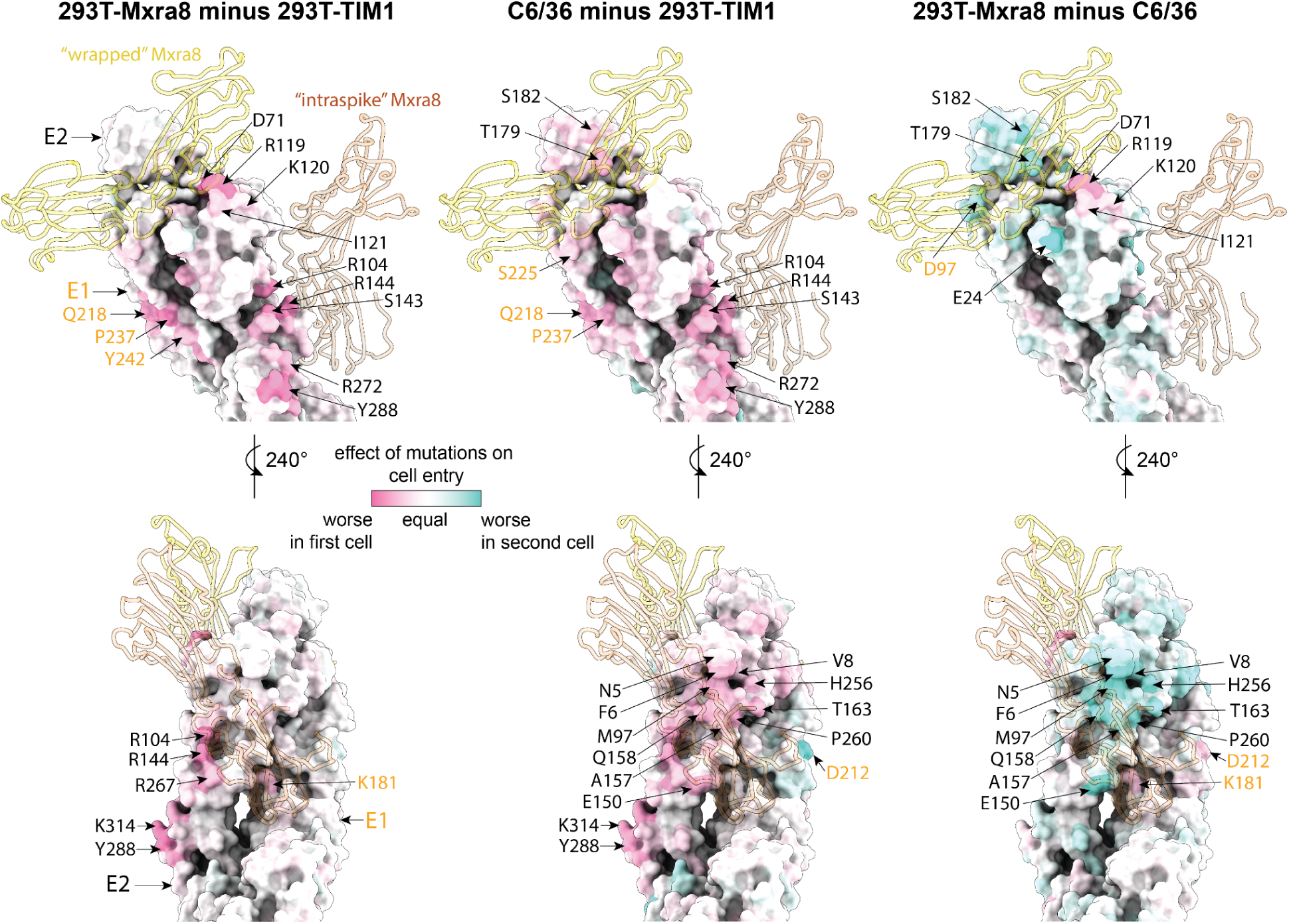
Structural locations of sites where mutations have different effects on entry among 293T-MXRA8, 293T-TIM1 and C6/36 cells. A single E2-E1 monomer from a cryo-EM structure of the full trimer (PDB: 6NK7) colored by the difference in mean effects of mutations at each site on entry between first and second cells. For instance, in the 293T-MXRA8 minus 293T-TIM1 plot (leftmost), red indicates sites where mutations are more deleterious in 293T-MXRA8 than 293T-TIM1 cells. MXRA8 is shown in in a thin ribbon in both the wrapped binding mode (gold) and the intraspike binding mode (light brown), with the MXRA8 in the intraspike binding mode modeled from another E2-E1 monomer in the cryo-EM structure of the full trimer (PDB: 6NK6) using ChimeraX matchmaker^98^. E2 sites (black text) and E1 sites (orange text) where mutations have large differences between different cell lines are labelled. E3 is removed from the structures to enable better visualization of the relevant E2 surfaces. See https://dms-vep.org/CHIKV-181-25-E-DMS/structures.html for interactive visualizations of these data on the protein structure.

We hypothesize that sites where mutations are more deleterious in C6/36 than 293T-TIM1 cells might similarly be involved in binding to the unknown mosquito receptor. Most sites where mutations impair entry in C6/36 cells more than 293T-TIM1 cells are in a large continuous surface on E2 (**Fig. 5**). Some sites where mutations are more deleterious in C6/36 cells than 293T-TIM1 cells are also more deleterious in 293T-MXRA8 than 293T-TIM1 cells and are near the MXRA8 binding interfaces (**Fig. 4c-d, 5**), suggesting there may be overlap of the surfaces used to bind MXRA8 and the mosquito receptor. However, there are some sites where mutations impair entry more in C6/36 than 293T-MXRA8 cells and vice versa (**Fig. 4b-d, 5**, **Extended Data Fig. 9**). For instance, mutations at sites 24, 95, 157 and 158 impair entry into C6/36 more than 293T-MXRA8 cells. Conversely, mutations at sites 71 and 119-121 impair entry in 293T-MXRA8 more than C6/36 cells. Note also that there are several sites in E1 (e.g., 210, 212, 386) where mutations are deleterious in both 293T-MXRA8 and 293T-TIM1 cells but tolerated in C6/36 cells (**Fig. 4b**); we hypothesize these sites may be important for some MXRA8-independent aspect of human cell entry.

To explore how differences in mutational constraints on the envelope proteins in human versus mosquito cells shape CHIKV evolution in nature, we examined the experimentally measured effects on entry in each cell line for all mutations in natural CHIKV envelope sequences relative to the 181/25 strain envelope used in our deep mutational scanning. Mutations observed in natural sequences were generally well tolerated for cell entry in all three cell lines in our deep mutational scanning, with naturally occurring mutations typically having effects near zero even though most mutations have negative effects on cell entry (**Extended Data Fig. 8a**). To test if natural selection for maintaining entry in both mosquito and human cells imposes additional constraint on natural evolution, we stratified all possible mutations by whether they are tolerated in both 293T-MXRA8 and C6/36 cells and found that mutations tolerated in both cells were more frequently observed in natural sequences than mutations tolerated in only one or neither cell type (**Extended Data Fig. 8b**). These results are consistent with natural selection during CHIKV evolution favoring mutations that maintain good envelope-mediated cell entry in both human and mosquito cells.

### Envelope protein mutants with decreased infectivity only in mosquito or MXRA8-expressing human cells

The above deep mutational scanning identified mutations that differentially affect pseudovirus entry in 293T-MXRA8 versus C6/36 cells (**Extended Data Fig. 9**). To validate these findings, we chose several mutations measured in the deep mutational scanning to impair entry only in 293T-MXRA8 cells or C6/36 cells and generated alphavirus RVPs expressing these mutations individually or in double-mutant combinations. We then measured the titers of these RVPs in 293T-MXRA8, 293T-TIM1 and C6/36 cells. Consistent with the deep mutational scanning, all mutants had titers similar to RVPs with unmutated envelope proteins in 293T-TIM1 cells (**Fig. 6a**). Also consistent with the deep mutational scanning, RVPs carrying the mutations expected to impair entry in 293T-MXRA8 cells (E2 mutations R119K, K120D and I121E) had over 10-fold reduced titers in those cells, but no major change in titers in C6/36 cells (**Fig. 6a**). Similarly, RVPs carrying the individual mutations expected to impair entry in C6/36 cells (E2 mutations A157S, Q158T and Q158V) had substantially reduced titers in those cells, but no major change in titers in 293T-MXRA8 cells. Combining mutations amplified the effect: double mutants expected to reduce entry in 293T-MXRA8 cells had over 100-fold reduced RVP titers in those cells but not C6/36 cells, and vice versa (**Fig. 6a**). To confirm that these effects were not specific to C6/36 and 293T-MXRA8 cells, we also measured the titers of all the RVPs in Aag2 cells (derived from *Aedes aegypti* mosquitoes^69,70^) and a subset of the RVPs in Huh7 and HFF-1 cells (human cells that naturally express MXRA8^15^). The titers in Aag2 cells mirrored those in C6/36 cells, whereas the titers in Huh7 and HFF-1 cells showed changes in the same direction as those in 293T-MXRA8 cells but with smaller magnitudes (**Fig. 6a**), possibly because the levels of MXRA8 or other attachment factors differ among these cell lines. Overall, these results suggest that our deep mutational scanning has identified mutations that generally reduce entry in either human cells expressing MXRA8 or mosquito cells.

**Figure 6.**
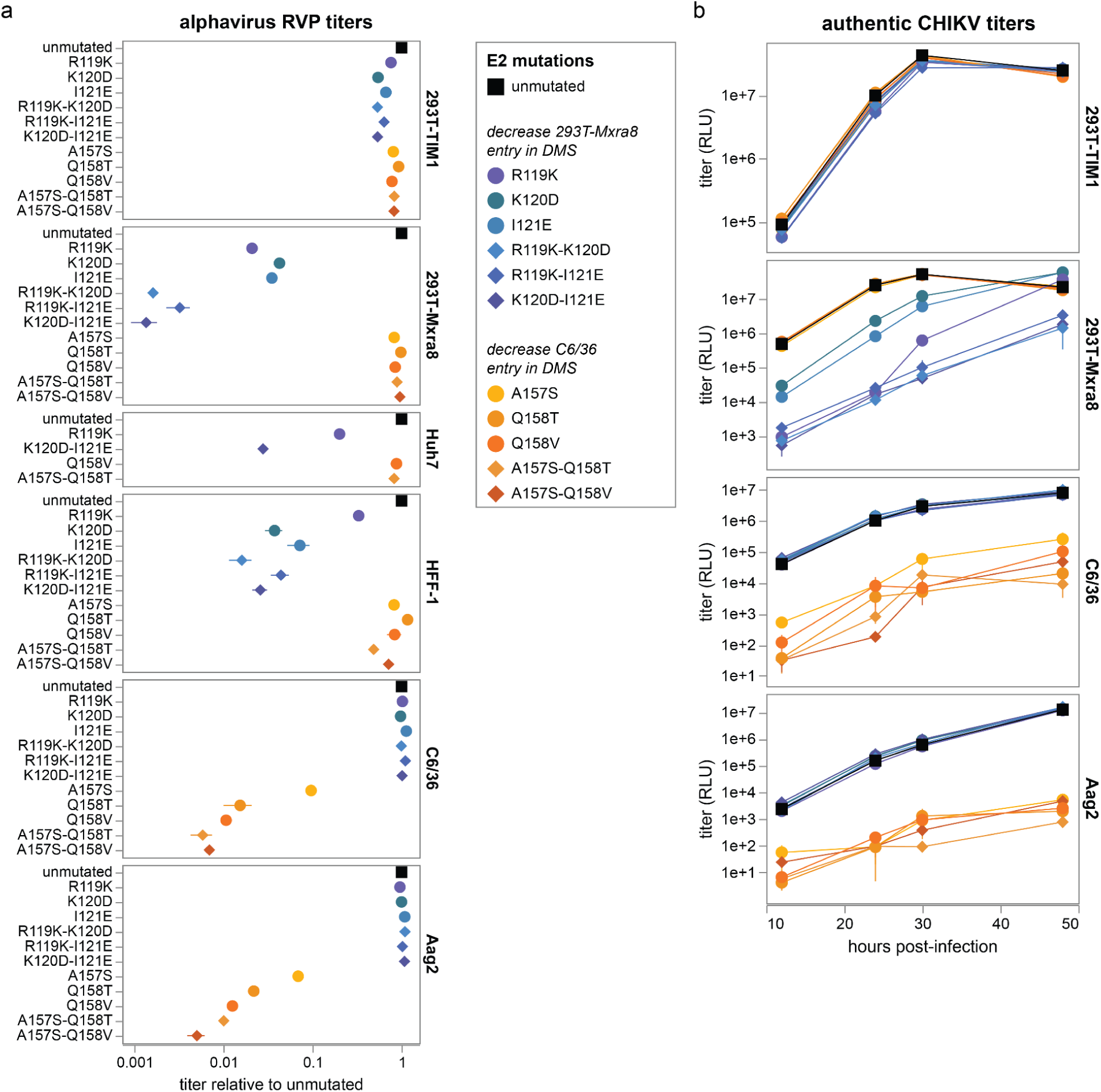
Mutants with reduced infectivity in mosquito or MXRA8-expressing human cells. (**a**) Titers of alphavirus RVPs carrying the indicated single or double mutation in E2 in different cell lines. Mutations were chosen because the pseudovirus deep mutational scanning indicated that they specifically impaired cell entry in either 293T-MXRA8 cells (mutations shown in various shades of blue) or C6/36 cells (various shades of orange). Note that Huh7 and HFF-1 are human cells that endogenously express MXRA8^15^; Aag2 are derived from *Aedes aegypti* mosquitoes^69,70^. Titers were quantified by the relative light units (RLU) produced by a luciferase reporter in the viral genome at 24 hours post infection and normalized to the titer in that cell line of RVPs with the unmutated envelope proteins. (**b**) Titers over time of replicative CHIKV (181/25 strain) in different cell lines. The viruses carry the indicated single or double mutations in E2. Titers were quantified by the relative light units (RLU) produced by a luciferase reporter in the viral genome at 12, 24, 30 and 48 hours after infection.

After demonstrating cell-specific loss of entry of the mutants in pseudovirus and alphavirus RVPs, we engineered the mutations into fully infectious CHIKV 181/25. These experiments do not constitute gain of function as the results for pseudovirus and RVPs establish that the mutations are loss of function in one cell type and neutral in the other; additionally the 181/25 strain of CHIKV is an attenuated strain^39,40^. As expected, viruses carrying the 293T-MXRA8 loss-of-entry mutations had reduced growth in those cells but wildtype-like growth in 293T-TIM1, C6/36, and Aag2 cells **(Fig. 6b**). Similarly, viruses carrying the C6/36 loss-of-entry mutations had reduced growth in C6/36 and Aag2 cells, but wildtype-like growth in 293T-MXRA8 and 293T-TIM1 cells **(Fig. 6b**). At early to intermediate timepoints, the cell-specific decrease of titers exceeded 100-fold for the 293T-MXRA8 loss-of-entry double mutants, and 10-fold for the mosquito-cell specific loss-of-entry mutants. These results show that pseudovirus deep mutational scanning can be used to design viruses with reduced tropism for either mosquito or MXRA8-expressing human cells. For the single and double mutants tested here, there remains some residual infectivity in the cell line for which tropism was reduced; however, it may be possible to further reduce this infectivity via additional mutations that the pseudovirus deep mutational scanning measures to impair cell-specific entry.

To confirm that the impaired entry of the species-specific loss-of-function mutations in MXRA8-expressing human or mosquito cells is because of reduced cell binding, we purified non-infectious CHIKV virus-like particles (VLPs) (**Extended Data Fig. 10a**) and measured their binding to 293T-MXRA8 or C6/36 cells. For these experiments, we generated non-infectious VLPs using the envelope proteins from the 37997 strain rather than 181/25 because the yield of 181/25 VLPs was too low, a limitation that has also been reported for the LR2006-OPY1 strain^71^. The E2 K120D-I121E double mutant had substantially reduced VLP binding to 293T-MXRA8 cells but still bound well to C6/36 cells, whereas the E2 V157S-Q158V double mutant had no VLP binding to C6/36 but still bound well to 293T-MXRA8 cells (**Extended Data Fig. 10b**). These results confirm that the species-specific loss of entry of these mutants is due to the mutations impairing binding to MXRA8 or the mosquito receptor, respectively.

## DISCUSSION

We used pseudovirus-based deep mutational scanning to comprehensively measure how mutations in the CHIKV envelope proteins affect entry in MXRA8-expressing human cells, TIM1-expressing human cells, and mosquito cells. While one prior study reported deep mutational scanning of CHIKV E3 and E2 in a single cell line^72^, our work is the first to enable systematic comparison of mutation effects across different cell lines with different receptors. The resulting maps of mutational effects elucidate sequence-function relationships, and inform efforts to engineer the envelope proteins as vaccine immunogens by identifying constrained epitopes. We also measured how envelope protein mutations affected binding to the MXRA8 receptor as assessed via pseudovirus neutralization by soluble MXRA8. Our identification of mutations that increase MXRA8 binding could aid complementary efforts to engineer soluble MXRA8 as an antiviral by making it bind more strongly to CHIKV^73^.

Our deep mutational scanning revealed mutations that specifically impair CHIKV entry in either MXRA8-expressing human cells or mosquito cells. These mutations shed light on determinants of CHIKV’s dual host tropism. CHIKV is one of several alphaviruses that naturally infect both vertebrate and invertebrate hosts. Vertebrate receptors have been identified for several of these dual-host alphaviruses including CHIKV, although the invertebrate receptors are less well characterized^15,53,54,74–76^. Although TIM1 and heparan sulfate promote CHIKV infection in mammalian cells^21,23,24^, they are unlikely to be the receptor(s) in mosquito cells. TIM1 orthologs have not been identified in mosquitoes^18^, and our deep mutational scanning shows that mosquito cell entry is impaired by mutations that do not impact entry in TIM1-expressing human cells. We could not detect heparan sulfate expression in the mosquito cells used in our experiments. Additionally, our deep mutational scanning shows that entry in C6/36 cells is improved by mutation E2 R82G, even though this mutation has previously been shown to impair heparan-sulfate usage by the CHIKV 181/25 strain in mammalian cells^22,77^. Work by others has also shown that soluble heparan sulfate inhibits CHIKV infection of some mammalian cells but not mosquito cells^18^. All this evidence suggests that entry in mosquito cells is not mediated by TIM1 or heparan sulfate; instead, our mapping of mutations that specifically impair entry in mosquito cells suggests that a broad continuous surface on E2 plays a key role in enabling infection of mosquito cells, possibly via binding to mosquito receptor(s). Knowledge of these mutations could inform future efforts to identify the mosquito receptor(s), such as via immunoprecipitation experiments using virions that can and cannot enter mosquito cells^78,79^.

Indeed, while this manuscript was under review, the Abraham laboratory posted a preprint reporting that the lachesin protein is a receptor for CHIKV in mosquito cells^80^. They used structural methods to show that lachesin interacts with E2 via a molecular surface very similar to the one identified in our deep mutational scanning measurements of sites where mutations specifically impair entry on mosquito cells. They further showed that the E2 S157-V158 variant described here abolishes lachesin binding^81^.

After using the deep mutational scanning to identify mutations that affected mosquito versus human cell entry in the context of non-replicative pseudoviruses, we designed loss-of-tropism mutants of the infectious 181/25 vaccine strain of CHIKV that were impaired in their ability to enter either mosquito cells or MXRA8-expressing human cells. This approach has potential applications to vaccine design if it can identify CHIKV mutants that have loss of function in actual mosquito infections as well as the mosquito cell lines used in our study. In particular, CHIKV with tropism restricted to cells from just a single species would be unable to undergo the full transmission cycle, and also likely attenuated in either mosquitoes or humans. Indeed, vaccination with the 181/25 strain caused transient arthralgia in some vaccine recipients^82^; it is possible such side effects could be reduced if the vaccine virus was unable to infect MXRA8-expressing human cells, as MXRA8-deficient mice showed reduced CHIKV-induced joint swelling^16^. Additionally, sites that are highly constrained for entry in either human or mosquito cells are promising targets for antibody development, since they represent functionally important regions where viral escape may be more restricted.

Our study has several limitations. First, our deep mutational scanning used a pseudovirus system based on lentiviral particles, which may not fully capture the entry dynamics or structural context of authentic CHIKV—although we did validate key mutations in alphavirus reporter particles or replicative CHIKV. Second, nearly all experiments used envelope proteins from the 181/25 strain, a live-attenuated strain that differs from its parental strain by four mutations in the envelope proteins^40^ that may impact protein behavior or cell binding^22^. Third, we measured entry in cell lines that may differ in their expression of potential receptor proteins relative to the cells infected by the virus *in vivo* in humans or mosquitoes, so future studies using *in vivo* models will be needed to validate that the mutations identified also attenuate actual virus in mammals or mosquitoes. Finally, we characterized the effects of individual mutations in a single genetic background, but in some cases the impacts of these mutations could differ in envelope proteins from different CHIKV strains due to epistasis^83–85^.

Despite these limitations, our study demonstrates how deep mutational scanning can comprehensively define the host-specific determinants of cell entry for a virus that naturally has tropism for species from two different phyla. In addition to providing clues about receptor usage and enabling the engineering of loss-of-tropism viral mutants, knowledge of these determinants can inform future analyses of how CHIKV evolution is shaped by selection to maintain the ability to infect cells from diverse species.

## ACKNOWLEDGMENTS

We thank Raul Andino (University of California San Francisco) for kindly providing Aag2 cells, Tamanash Bhattacharya and Harmit Malik (Fred Hutch) for C6/36 cells, James Crowe (Vanderbilt University) for sharing the 4J21, 4N12, CHKV-24, and CHKV-35 antibodies, Roland Strong (Fred Hutch) for providing access to laboratory equipment and Scott Weaver (University of Texas Medical Branch) for the CHIKV 181/25 infectious cDNA clone. We thank Bernadeta Dadonaite and Wenlin Ren for technical advice and feedback. The AI agent Claude was used for some of the data analysis and manuscript proofreading. The cartoon diagrams of lentiviruses and cells in Extended Data Figures 1, 2 and 3 were created with BioRender.com. This work was supported by the NIH/NIAID under the FLARE ReVAMPP (U19AI181960 to MSD, DHF, and JDB), R01AI143673 (to MSD and DHF), NIAID contract 75N93022C00035 (to DHF) and R01AI141707 (to JDB). We acknowledge support from the Genomics & Bioinformatics Shared Resource (RRID:SCR_022606) and Flow Cytometry Shared Resource (RRID:SCR_022613) of the Fred Hutch/University of Washington/Seattle Children’s Cancer Consortium (P30 CA015704), and Fred Hutch Scientific Computing (NIH grants S10-OD-020069 and S10-OD-028685). JDB is an Investigator of the Howard Hughes Medical Institute. This manuscript is the result of funding in whole or in part by the National Institutes of Health (NIH). It is subject to the NIH Public Access Policy. Through acceptance of this federal funding, NIH has been given a right to make this manuscript publicly available in PubMed Central upon the Official Date of Publication, as defined by NIH.

## METHODS

### Data and code availability

All processed data, interactive visualizations, and sequencing counts for each experiment are publicly available on GitHub. See the homepage at https://dms-vep.org/CHIKV-181-25-E-DMS/ for interactive visualizations, links to CSV files with the numerical data, and an appendix with details from the notebooks used to analyze the data. See the GitHub repository at https://github.com/dms-vep/CHIKV-181-25-E-DMS for all computer code and processed data.

### Biosafety

All experiments with pseudoviruses, RVPs, and virus were performed at biosafety-level-2. The deep mutational scanning used pseudoviruses (lentiviral pseudotyped particles) that encode no viral proteins other than the CHIKV envelope proteins, and only undergo a single round of cell entry. Some validation experiments were performed using alphavirus reporter virus particles (RVPs) that encode the replicase proteins from the attenuated VEEV TC-83 strain with the structural proteins provided in trans on two separate plasmids and not encoded in the viral genome; these RVPs undergo a single round of cell entry. A few experiments were performed with the fully infectious CHIKV 181/25 strain; this is an attenuated vaccine strain that has been safety-tested in humans and approved for use at biosafety-level-2^39,40,82^. Mutations were introduced into CHIKV 181/25 only after they had been confirmed with both pseudoviruses and RVPs to confer a loss-of-function (reduced cell tropism) phenotype.

### Cell lines

The following cell lines were used: 293T (ATCC, CRL-3216), 293T-rtTA (from Dadonaite et al.^37^), 293T-TIM1 (from Simonich et al.^64^), Huh7 (JCRB, JCRB0403), C6/36 (from Bhattacharya et al.^86^), Aag2 (from Tassetto et al.^87^), CHO-pgsA745 (ATCC, CRL-2242), HFF-1 (ATCC, SCRC-1041) and Vero E6 (ATCC, CRL-1586). 293T, Huh7, Vero E6, HFF-1 and 293T derivative cells were cultured in D10 media (Dulbecco’s Modified Eagle Medium supplemented with 10% heat-inactivated fetal bovine serum, 2 mM l-glutamine, 100 U/mL penicillin, and 100 μg/mL streptomycin) at 37°C with 5% CO2. C6/36 and Aag2 cells were cultured in M10 (Minimum Essential Medium supplemented with 10% heat-inactivated fetal bovine serum, 2 mM l-glutamine, 1×MEM non-essential amino acids solution, 100 U/mL penicillin, and 100 μg/mL streptomycin) at 28°C with 5% CO2. CHO-pgsA745 cells were cultured in F10 (Ham’s F-12K (Kaighn’s) Medium supplemented with 10% heat-inactivated fetal bovine serum, 2 mM L-glutamine, 100 U/mL penicillin, and 100 μg/mL streptomycin) at 37°C with 5% CO2. To inhibit rtTA activation, 293T-rtTA cells were maintained in tet-free D10, which is made with tetracycline-negative fetal bovine serum (Gemini Bio, 100-800).

### Plasmids and primers

Plasmid maps can be found at:

https://github.com/dms-vep/CHIKV-181-25-E-DMS/tree/main/data/supplemental_data/plasmids

Primer sequences can be found at:

https://github.com/dms-vep/CHIKV-181-25-E-DMS/blob/main/data/supplemental_data/primers.csv

### 293T-MXRA8 cells

Human MXRA8 (NP_001269511) followed by internal ribosome entry site (IRES) and mCherry was cloned into the pHAGE2 lentivirus vector. Lentivirus pseudotyped with VSV-G was rescued from 293T cells and used to transduce 293T cells. After picking and expanding a single 293T clone, immunostaining was performed to confirm the expression of MXRA8. For immunostaining, 293T and 293T-MXRA8 cells were stained with anti-MXRA8 primary antibody (MBL International, W040-3) diluted in 1% BSA at 1:100 for 1 hour on ice and 488-conjugated goat anti-mouse IgG, IgM secondary antibody (Invitrogen, A-10680) diluted in 1% BSA at 1:1000 for 1 hour on ice. Cells were analyzed by flow cytometry to confirm MXRA8 expression.

### Design of envelope proteins deep mutational scanning libraries

The libraries were designed in the background of the CHIKV 181/25 strain (MW473668.1). Envelope proteins (E3-E2-6K-E1) were codon optimized via GeneArt tool, as this optimization increased pseudovirus titer. The plasmid map for the lentiviral backbone with codon-optimized envelope proteins sequence is at: https://github.com/dms-vep/CHIKV-181-25-E-DMS/blob/main/data/supplemental_data/plasmids/4581_V5LP_pH2rU3_ForInd-Extgag_181-25_E_CO4_GA_CMV_ZsGT2APurR.gb. As envelope proteins are a large polyprotein, we mutagenized the gene sequence in two halves (E3-E2 libraries and 6K-E1 libraries), although we subsequently combined the libraries for many of the experiments reported here. We ordered the libraries for each of these two halves in the form of linear dsDNA from GenScript Biotech as a site-saturation variant library containing all possible single amino acid mutations in E3 and E2 (E3E2) and another site-saturation variant library containing all possible single amino acid mutations in 6K and E1 (6KE1). In addition to including codons for all 19 amino-acid mutations at each site, we included 20 stop codons located at alternating positions from the second residue of E3 and 6K as negative controls for cell entry measurements. Cell entry measurements were conducted separately using the E3E2 and 6KE1 libraries, and the results were subsequently combined computationally. MXRA8 binding measurements were performed using the combined E3E2–6KE1 libraries. The final quality control report for the library is available at: https://github.com/dms-vep/CHIKV-181-25-E-DMS/tree/main/data/supplemental_data/GenScript_QC_report

### Cloning of the envelope proteins mutant library into lentiviral backbone plasmid

The linear dsDNA encoding the mutagenized E3-E2 or 6K-E1 region of the envelope proteins was cloned into a lentiviral backbone plasmid for pseudovirus deep mutational scanning as described previously^37^. Barcoding PCR that appends a barcode consisting of 16 random nucleotides downstream of the envelope protein gene stop codon was performed by adding 20 ng (1 μL) of the GenScript library DNA, 1.5 μL of ForInd_AddBC_2 primer (10 μM), 1.5 μL of 5’for_lib_bcing primer (10 μM), 21 μL of nuclease free water, and 25 μL of KOD Hot Start Master Mix (ThermoFisher, 71842-4). The PCR cycling conditions were:

1. 95℃ for 2 min
2. 95℃ for 20 sec
3. 55.5℃ for 20 sec (cooling at 0.5 °C/sec)
4. 70℃ for 2 min
5. Return to Step 2, 9 cycles
6. 4℃ hold

To have biological replicates, the barcoding was performed in two independent reactions for each library, producing two separate PCR products with unique barcodes (A and B). So, we had two E3E2 libraries (E3E2_A and E3E2_B) and two 6KE1 libraries (6KE1_A and 6KE1_B) eventually. All subsequent cloning and virus rescue steps were conducted separately for each library. The lentiviral backbone was prepared by digesting plasmid 4016_V5LP_pHrU3_ForInd-Extgag_mcherry (https://github.com/dms-vep/CHIKV-181-25-E-DMS/blob/main/data/supplemental_data/plasmids/4016_V5LP_pHrU3_ForInd-Extgag_mcherry.gb) with MluI-HF and XbaI for 3 hours at 37°C. The PCR products and digested vector backbone were run on a 1% agarose gel, and bands of the expected size were excised and cleaned up with a NucleoSpin Gel and PCR Clean-up kit (Macherey-Nagel, 740609.5) followed by additional cleanup with Ampure XP beads (Beckman Coulter, A63881). Barcoded envelope protein libraries were cloned into the lentiviral backbone using the NEBuilder HiFi DNA Assembly Kit (NEB, E2621) with a 2:1 insert to vector molar ratio in a 1-hour reaction at 50℃. The HIFI reactions were purified with Ampure XP beads and eluted in nuclease free water, then transformed into 10-beta electrocompetent cells (NEB, C3020K) with a BioRad MicroPulser Electroporator (BioRad, 1652100), shocking at 2 kV for 5 milliseconds. 15 electroporation reactions were performed for each library. Following a 1-hour recovery at 37°C, transformed cells were pooled and cultured in 300 mL LB medium supplemented with ampicillin for 16 hours at 37°C with shaking. Plasmids were extracted using the QIAGEN HiSpeed Plasmid Maxi Kit (QIAGEN, 12662). The total number of colonies for E3E2_A, E3E2_B, 6KE1_A and 6KE1_B were 3.0e6, 2.2e6, 2.6e6 and 2.8e6 CFUs respectively. A large number of separate colonies are necessary to prevent bottlenecking of our library diversity and to ensure that any recombination of the diploid lentiviral genomes during integration of the library into cells does not associate the same barcode with different variants (to avoid this, the number of colonies should be much greater than the ultimate size of the library integrated into the cells).

### Creation of cell-stored deep mutational scanning libraries

To perform deep mutational scanning, we need genotype-phenotype linked pseudoviruses. The rationale for this is described in prior work^37^ and **Extended Data Fig. 1b**. Following a previously described protocol^37^, we generated cell-stored deep mutational scanning libraries where each cell was integrated with a single copy of a barcoded envelope protein mutant enabling rescue of genotype-phenotype linked pseudoviruses. 10cm dishes were seeded with ∼10^7^ 293T cells. On the next day, each dish was transfected with 6 μg of plasmid library encoding the lentiviral backbone with barcoded envelope protein mutants, 1.5 μg of each lentiviral helper plasmid (26_HDM_Hgpm2 (Addgene, 204152), 27_HDM_tat1b (Addgene, 204154), and 28_pRC_CMV_Rev1b (Addgene, 204153)) and 1.5 μg of plasmid expressing VSV-G (29_HDM_VSV_G (Addgene, 204156)) using BioT transfection reagent (Bioland Scientific, B01-02). Two 10cm dishes were used for each library, resulting in ∼20 mL of VSV-G pseudotyped library viruses. At 48 hours post transfection, viruses were filtered through a 0.45 μm syringe filter (Corning, 431220), pooled, aliquoted and stored at −80°C. An aliquot of these viruses was used to infect 293T-rtTA cells, and the titer was determined by measuring the percentage of ZsGreen positive cells via flow cytometry.

After titering, the VSV-G pseudotyped library viruses were used to infect 293T-rtTA cells (as described previously^37^, these cells produce high viral titers) at 0.7% multiplicity of infection (MOI) to ensure most cells are infected by at most one virion. The MOI was confirmed by measuring the percentage of ZsGreen positive cells via flow cytometry at 48 hours post infection. Based on the measured ZsGreen positive rate and cell number during infection, cells were pooled so that each library would contain about 60,000 infected cells. We chose this number to ensure each mutant is associated with multiple barcodes, which increases measurement accuracy (for ∼10,000 mutants, each mutant would have ∼6 barcodes), while being low enough to ensure all variants are measured during the selection conditions. The libraries ended up being close to this target: 65,426 for E3E2_A, 62,495 for E3E2_B, 69,359 for 6KE1_A and 62,446 for 6KE1_B. Integrated cells were selected by 0.75 μg/mL of puromycin for 1 week. Cells were expanded in tet-free D10 for 48 hours and then frozen down in the gas phase of liquid nitrogen in aliquots of 2e7 cells for long-term storage.

### Generation of VSV-G and envelope protein expressing pseudovirus libraries

To generate pseudoviruses expressing envelope proteins from integrated cell lines, 1.5 x 10^8^ 293T-rTTA cells with the integrated libraries were seeded into 5-layer cell culture flasks (Corning, 353144) in tetracycline-free D10 medium supplemented with 0.1 μg/mL doxycycline to induce envelope protein expression from the integrated genome. The following day, each flask was transfected with 50 μg of each helper plasmid (26_HDM_Hgpm2, 27_HDM_tat1b, and 28_pRC_CMV_Rev1b) using BioT transfection reagent (according to the manufacturer’s instructions). At 48 hours post-transfection, the culture supernatant was harvested and filtered through a 0.45 μm SFCA Nalgene 500 mL Rapid-Flow filter unit (ThermoFisher, 09-740-44B). The filtered supernatant was concentrated by ultracentrifugation at 100,000 × g for 1 hour at 4 °C. Viral pellets were resuspended in D10 medium to achieve a 50-fold concentration. Aliquots of 1.5 mL were prepared and stored at −80 °C for downstream deep mutational scanning experiments.

To rescue pseudoviruses expressing VSV-G from integrated cell lines,, 2.4 x 10^7^ cells were seeded into 15 cm dishes in tetracycline-free D10 medium. The following day, each dish was transfected with 7.5 μg of each helper plasmid (26_HDM_Hgpm2, 27_HDM_tat1b, and 28_pRC_CMV_Rev1b) as well as 7.5 μg of plasmid 29_HDM_VSV_G encoding the VSV-G. At 48 hours post-transfection, the supernatant was harvested and filtered through a 0.45 μm SFCA Nalgene 500 mL Rapid-Flow filter unit. The filtered supernatant was concentrated using Lenti-X Concentrator (Takara, 631232) at a 1:3 virus-to-concentrator volume ratio, incubated overnight at 4 °C, and centrifuged at 1,500 × g for 45 minutes at 4 °C. Viral pellets were resuspended in D10 medium to achieve a 12-fold concentration. Aliquots of concentrated VSV-G-expressing pseudoviruses were stored at −80 °C for use in barcode-mutation linkage and cell entry deep mutational scanning experiments.

The titer of pseudoviruses in transduction units (TU) per ml was determined by measuring the percentage of ZsGreen positive cells via flow cytometry. Typical titers for our experiments were ∼1e5 and ∼3e5 TU/ml for pseudovirus libraries generated from the integrated cells prior to concentration in 293T-MXRA8 and 293T-TIM1 cells, and ∼5e6 and ∼1.5e7 TU/ml in 293T-MXRA8 and 293T-TIM1 after concentration. Note that we can not measure titers of pseudoviruses in TU in C6/36 cells since the lentivirus does not transcribe the integrated reporter genes in these cells, but for our titer estimations assumed the infectivity of pseudoviruses in C6/36 was similar to 293T-MXRA8 cells.

### Long-read PacBio sequencing to link mutations to barcodes

As described previously^37^, the barcodes must be linked to the envelope proteins variants after integration of the libraries into the cells that store them, as this integration can scramble the linkage of barcodes to gene variants relative to the plasmid library due to recombination of the co-packaged pseudodiploid lentiviral genomes. To perform the linkage, 293T cells were seeded at a density of 3 x 10^6^ cells per well in poly-L-lysine-coated 6-well plates. The following day, approximately 1.5 x 10^7^ transducing units (TU) of VSV-G expressing pseudoviruses that were generated from the cell-stored libraries were used to infect each of 3 wells. At 12 hours post-infection, non-integrated reverse-transcribed lentiviral genomes^88,89^ were extracted from the infected cells using the QIAprep Spin Miniprep Kit (Qiagen, 27104) according to the manufacturer’s protocol, which enables recovery of episomal DNA. The amplicon for PacBio sequencing was prepared following a protocol originally described in Dadonaite et al^37^. Miniprepped DNA was split into two parallel PCR reactions to assess strand exchange events by incorporating distinct single-nucleotide identifiers at the 5′ and 3′ ends of the amplicon. Both reactions used the same PCR conditions: 20 μL KOD Hot Start Master Mix, 18 μL of miniprepped DNA, and 1 μL each of 10 μM primers. Reaction 1: 5_PacBio_G and 3_PacBio_C. Reaction 2: 5_PacBio_C and 3_PacBio_G. Round 1 PCR cycling conditions:

1. 95 °C for 2 min
2. 95 °C for 20 sec
3. 70 °C for 1 sec
4. 60 °C for 10 sec (cooling at 0.5 °C/sec)
5. 70 °C for 70 sec
6. Repeat steps 2–5 for 7 cycles
7. 70 °C for 70 sec
8. Hold at 4 °C

Round 1 PCR products were purified using 50 μL of Ampure XP beads and eluted in 35 μL of nuclease free water. Equal volumes of the two reactions were pooled for each library. Round 2 PCR setup: each reaction contained 25 μL of KOD Hot Start Master Mix, 21 μL of pooled round 1 product, and 2 μL each of 10 μM 5_PacBio_Rnd2 and 3_PacBio_Rnd2 primers. Round 2 PCR cycling conditions:

1. 95 °C for 2 min
2. 95 °C for 20 sec
3. 70 °C for 1 sec
4. 60 °C for 10 sec (cooling at 0.5 °C/sec)
5. 70 °C for 70 sec
6. Repeat steps 2–5 for 10 cycles
7. 70 °C for 70 sec
8. Hold at 4 °C

Both these PCRs use a limited number of cycles to avoid PCR strand exchange that would scramble the barcodes relative to the initial template. Final PCR products were purified using 50 μL of Ampure XP beads and eluted in 40 μL of nuclease-free water. PCR products from each library were combined, and amplicon size was verified using TapeStation. Libraries were sequenced using circular consensus sequencing on a PacBio Sequel IIe sequencer. For details on downstream data processing, refer to the **PacBio sequencing analysis to link barcodes to envelope-protein mutants** section.

### Deep mutational scanning experiments to measure effects of mutations on cell entry

To measure the effects of mutations on cell entry, we followed the approach described by Dadonaite et al.^37^. Briefly, separate wells of cells were infected with (i) a pseudovirus library expressing the envelope proteins and (ii) a pseudovirus library expressing VSV-G. The VSV-G pseudovirus served as a baseline control to determine library composition, as VSV-G mediates cell entry independently of the envelope proteins. For each infection, 2 x 10^6^ 293T-MXRA8, 293T-TIM1 cells in D10 medium and C6/36 in M10 medium were seeded per well of a 6-well plate. The following day, cells from three wells were infected with ∼2e6 TU of envelope protein pseudovirus library or ∼8e6 TU of VSV-G pseudovirus library. Note that C6/36 cells in M10 media were moved from 28°C to 37 °C right after infection to increase the yield of non-integrated reverse-transcribed lentiviral genomes, as reverse transcriptase activity is higher at 37 °C than 28 °C^90^. (Note that although the lentivirus does not transcribe the integrated reporter genes in C6/36 cells, it still produces the non-integrated reverse-transcribed lentiviral genomes that we sequence.) At 12 hours post infection, non-integrated reverse-transcribed lentiviral genomes^88,89^ were recovered by miniprepping. Amplicons for Illumina sequencing were prepared in two PCR rounds. In Round 1, the Illumina TruSeq Read 1 and Read 2 sequences were appended. Each 50 µL Round 1 reaction contained: 22 µL miniprepped DNA, 25 µL KOD Hot Start Master Mix, 1.5 µL Illumina_Rnd1_For primer (10 µM), and 1.5 µL Illumina_Rnd1_Rev3 primer (10 µM). Cycling conditions were:

1. 95 °C for 2 min
2. 95 °C for 20 sec
3. 70 °C for 1 sec
4. 58 °C for 10 sec (cooling at 0.5 °C/sec)
5. 70 °C for 20 sec
6. Repeat steps 2–5 for 27 cycles
7. 70 °C for 60 sec
8. Hold at 4 °C

Round 1 products were purified with 150 µL Ampure XP beads and eluted in 50 µL nuclease-free water. DNA concentrations were measured using a Qubit 4 Fluorometer (ThermoFisher, Q33238). Round 2 PCR attached sample indices for multiplexing. Each reaction contained 20 ng Round 1 product, 25 µL KOD Hot Start Master Mix, 2 µL of each Round 2 indexing primer (10 µM), and nuclease-free water to 50 µL total. Cycling conditions matched Round 1 except only 20 cycles were performed. DNA concentrations were again determined by Qubit 4. Round 2 products were pooled in equimolar amounts, resolved on a 1% agarose gel, and the 283 bp band was excised. DNA was purified using the NucleoSpin Gel and PCR Clean-up Kit, followed by Ampure XP bead purification. Libraries were diluted to 10 nM and sequenced on an Illumina NovaSeq X Plus system. For details on how sequencing counts were converted to mutation effects on cell entry, see the **Analysis to determine mutation effects on cell entry in pseudovirus deep mutational scanning** section.

### Generations and purification of soluble mouse and human MXRA8-Fc

Soluble mouse and human MXRA8 were generated and purified as previously described^15^. Briefly, cDNA fragments encoding the extracellular domain of mouse MXRA8 (residues 22–336; NM_024263) or human MXRA8 (residues 20–337; NM_032348.3), fused to mouse IgG2b–Fc, were synthesized and cloned into the pCDNA3.4 vector. Soluble MXRA8–Fc proteins were expressed in Expi293 cells (Thermo Fisher, A14527) at a density of 3 x 10^6^ cells/mL. For transfection, 200 µg plasmid DNA was diluted in Opti-MEM (Thermo Fisher, 31985070) and mixed with 640 µL ExpiFectamine 293 Reagent. After a 15-minute incubation, ExpiFectamine/DNA complexes were added to the cells. At 22 h post transfection, 1.2 mL ExpiFectamine 293 Transfection Enhancer 1 and 12 mL ExpiFectamine 293 Transfection Enhancer 2 were added. Four days post transfection, culture supernatants were collected and clarified by centrifugation at 30,000 × g for 15 min. Soluble MXRA8–Fc proteins were purified by protein A Sepharose 4B (Thermo Fisher, 101041) affinity chromatography. Eluted proteins were immediately neutralized, dialyzed into 20 mM HEPES, 150 mM NaCl (pH 7.5), filtered through a 0.22 µm membrane, and stored at −80 °C.

### Production and purification of CHIKV virus-like-particles (VLPs)

CHIKV VLPs used in the VLP cell-surface binding experiments shown in Extended Data Figure 10 were produced via transient transfection of plasmid DNA encoding C-E3-E2-6K-E1 (CHIKV strain 37997)^51^ into Expi293 cells. Because these VLPs do not encode genes they are incapable of replicating any genetic material, meaning that they are not infectious viruses and so are safe for study at biosafety level 2. For 1 L cell culture, the cells were transfected with a mixture of 1 mg CHIKV VLP plasmid and 3 mg polyethylenimine (PEI; Polysciences, 24765-2) at a cell density of 3 x 10^6^ cells/mL. VLPs were collected at 24 hours post transfection and purified using low-speed spin centrifugation^91^. VLPs were resuspended in TNE-V buffer (25mM Tris, 150mM NaCl, 2mM EDTA pH 8.0) and stored at 4 °C. VLPs were analyzed by electrophoresis on 4–20% Mini-PROTEAN TGX Stain-Free Gels (Bio-Rad, 17000435).

### Deep mutational scanning experiments to measure effects of mutations on MXRA8 binding

To measure the effects of mutations on MXRA8 binding, we followed the approaches described by Dadonaite et al.^57^ for soluble ACE2 and Larsen et al.^38^ Briefly, 2 x 10^6^ 293T-MXRA8 cells in D10 medium were seeded into each well of a 6-well plate. The following day, ∼2e6 TU of pooled envelope-protein pseudovirus library was mixed with ∼2e4 TU of VSV-G neutralization standard viruses^37^ (VSV-G pseudotyped viruses are not neutralized by soluble MXRA8-Fc) containing eight distinct, known barcodes. The mixed viruses were incubated with different concentrations of soluble mouse MXRA8-Fc (0, 6.4, 20, 60, 180, or 540 µg/mL) or soluble human MXRA8-Fc (0, 6.4, 20, 60, 120, or 240 µg/mL) for 1 hour at 37 °C. After incubation, the virus mixtures were used to infect 293T-MXRA8 cells. At 12 hours post infection, non-integrated reverse-transcribed lentiviral genomes were recovered by miniprepping, and amplicons for Illumina sequencing were prepared as described above. Our measurements of effects of mutations on human MXRA8 binding were more noisy than the measurements on mouse MXRA8 binding since human MXRA8 is less potent in neutralizing the pseudoviruses and we used a lower maximal concentration. Details on the computational conversion of sequencing counts to mutation effects on MXRA8 binding are provided in the section **Analysis to determine mutation effects on MXRA8 binding in pseudovirus deep mutational scanning**.

### Generation of alphavirus reporter virus particles (RVPs)

RVPs were generated as previously described^53^ with some modifications. Briefly, 293T cells were transfected with three plasmids using BioT transfection reagent. The first plasmid encoded a VEEV TC-83 replicon (https://github.com/dms-vep/CHIKV-181-25-E-DMS/blob/main/data/supplemental_data/plasmids/4996_VEEV-TC-83-Fluc2AEGFP.gb) in which the structural protein genes were replaced by firefly luciferase and EGFP, with expression driven by the Rous sarcoma virus (RSV) enhancer/promoter. The second and third plasmids encoded the CHIKV capsid from the LR2006 OPY1 strain (https://github.com/dms-vep/CHIKV-181-25-E-DMS/blob/main/data/supplemental_data/plasmids/5008_HDM_CHIKV_Capsid.gb) and CHIKV envelope proteins (E3-E2-6K-E1) from the 181/25 strain (https://github.com/dms-vep/CHIKV-181-25-E-DMS/blob/main/data/supplemental_data/plasmids/4762_HDM_181-25ECO4.gb), respectively, under control of the CMV promoter. Prior to transfection, 293T cells were seeded into 6-well plates at a density of 2 x 10^6^ cells per well. The following day, 1 μg plasmid encoding VEEV TC-83 replicon, 0.5 μg plasmid encoding CHIKV capsid and 0.5 μg plasmid encoding CHIKV envelope proteins were transfected using BioT. Supernatants were harvested at 48 hours post transfection, clarified by filtration through a 0.45 μm filter, and aliquoted for storage at −80 °C.

### Validations of deep mutational scanning measurements using reporter virus particles (RVPs)

RVPs produced using unmutated and the indicated mutant envelope proteins were generated by co-transfection of three plasmids into 293T cells as described above. For cell entry validations, 293T-MXRA8, 293T-TIM1, Huh7, HFF-1, C6/36, or Aag2 cells were seeded into blackwalled, 96-well plates (Greiner, 655090) at a density of 40,000 cells per well. The following day, cells were infected with 1 μL of wildtype or mutant RVPs. To validate deep mutational scanning measurements for soluble MXRA8-Fc neutralization, the same method was used, except RVPs were incubated with soluble MXRA8-Fc for one hour prior to adding to 293T-MXRA8 cells. Each plate contained cell-only and RVP-only control wells to correct for any background luciferase signal. Luciferase activity was measured at 24 hours post infection. For each measurement, 150 μL of supernatant was removed from each well, and 30 μL of Bright-Glo Luciferase Assay reagent (Promega, E2610) was added. Luminescence was immediately recorded using a Tecan Infinite M1000 plate reader. IC50 values were estimated by fitting neutralization curves using the package neutcurve (https://github.com/jbloomlab/neutcurve)^92^. Experiments with the human cells were performed at 37°C whereas experiments with the mosquito cells were performed at 28°C.

### Creation of replicative CHIKV expressing reporter proteins

An infectious cDNA clone of CHIKV 181/25^40^ (obtained from Scott Weaver, University of Texas Medical Branch) was modified to express firefly luciferase and EGFP reporters. Briefly, the SP6 promoter was replaced with the Rous sarcoma virus (RSV) enhancer/promoter, enabling virus rescue via plasmid transfection into 293T cells. Reporter genes were placed under the control of a duplicate subgenomic (SG) promoter, as previously described^93^. The full Genbank map of the plasmid encoding this replicative reporter CHIKV is available at: https://github.com/dms-vep/CHIKV-181-25-E-DMS/blob/main/data/supplemental_data/plasmids/5395_RSVP_181-25ic_Fluc2AEGFP_HDVR.gb. To produce virus from this infectious clone, 293T cells were seeded into 6-well plates at a density of 2 x 10^6^ cells per well. The following day, 2 μg plasmid was transfected using BioT. Supernatants were collected at 24 hours post transfection, clarified by filtration through a 0.45 μm filter, and aliquoted for storage at −80 °C.

### Replication-competent CHIKV growth kinetics assays

CHIKV wild-type and mutant viruses were generated in 293T cells via plasmid transfection. 293T-MXRA8, 293T-TIM1, C6/36, and Aag2 cells were seeded into 96-well plates at a density of 4 x 10^4^ cells per well. The following day, cells were infected with 1 μL of virus per well. Each plate contained cell-only and virus-only control wells to correct for any background luciferase signal. Luciferase activity was measured at 12, 24, 30, and 48 hours post infection. For each measurement, 150 μL of supernatant was removed from each well, and 30 μL of Bright-Glo Luciferase Assay reagent (Promega, E2610) was added. Luminescence was immediately recorded using a Tecan Infinite M1000 plate reader. Experiments with the human cells were performed at 37°C whereas experiments with the mosquito cells were performed at 28°C.

### Heparan sulfate staining

Cells were digested by 0.25% trypsin and washed with PBS. 10^6^ cells were stained with anti-heparan sulfate primary antibody (Amsbio, 370255-1) diluted in 1% BSA at 1:100 for 1 hour on ice and 488-conjugated goat anti-mouse IgG, IgM secondary antibody (Invitrogen, A-10680) diluted in 1% BSA at 1:1000 for 1 hour on ice. Cells were resuspended in 1% BSA and analyzed by flow cytometry.

### Cell-based binding assay

To measure the interaction between CHIKV VLPs and cells, 293T-MXRA8 cells were trypsinized with 0.25% trypsin, whereas C6/36 cells were detached by pipetting. 10^6^ cells were incubated with 20 μg VLPs for 1 hour on ice. After two washes with DPBS, cells were incubated with a mixture of mAbs (4J21^32^, 4N12^32^, CHKV-24^25^, and CHKV-35^25^; 2 μg/mL each) for 1 hour on ice and 2 μg/mL 488-conjugated goat anti-human IgG, IgM secondary antibody (Invitrogen, A-11013) for 1 hour on ice. After two washes with DPBS, cells were resuspended in 1% BSA and analyzed by flow cytometry. To measure the interaction between cell-expressed E proteins and MXRA8-Fc, we transfected 293T cells with a plasmid encoding the E proteins. Cells were detached by Cell Dissociation Buffer (Gibco, 13150016) at 24 hours post transfection. After a wash with DPBS, 10^5^ cells were incubated with varying concentrations of soluble mouse MXRA8-Fc for 1 hour on ice and 2 μg/mL 488-conjugated goat anti-human IgG, IgM secondary antibody for 1 hour on ice. After two washes with DPBS, cells were resuspended in 1% BSA and analyzed by flow cytometry.

### PacBio sequencing analysis to link barcodes to envelope-protein mutants

To link the barcodes encoded in the lentiviral genomes to the envelope protein variant on the same genome, the PacBio circular-consensus sequences (CCSs) were aligned to the expected sequence of the amplicons and processed using alignparse^94^. The empirical accuracy of the CCSs was ∼60% for high-quality CCSs with no out-of-frame indels, as assessed by determining how often CCSs with the same barcode were associated with the same envelope proteins gene variant. The imperfect empirical accuracy is due to some combination of sequencing errors, a small amount of strand-exchange during the PCR library preparation, reverse transcription errors and lentiviral recombination during library integration into cells leading the same barcode to be associated with multiple envelope proteins variants. To create a high-accuracy barcode-variant lookup table, we built consensus sequences using only barcodes with at least three CCSs that did not have any non-consensus mutations at appreciable frequency across CCSs. This analysis was all performed in the context of dms-vep-pipeline-3 (version 3.27.0, https://github.com/dms-vep/dms-vep-pipeline-3); the computational notebooks performing this analysis and a hyperlink to the final barcode-variant lookup table are at https://dms-vep.org/CHIKV-181-25-E-DMS/appendix.html in the subsection titled “Barcode to codon-variant lookup table.” Note that the analysis was performed separately for the E3-E2 and 6K-E1 for each library as well as a combined library (E3-E3-6K-E1) analysis that simply concatenated the CCSs for the two halves of the gene. Some downstream experiments were performed using the separate library halves with the results then combined whereas others were performed on pools of the two library paths, so the appropriate barcode-variant lookup table was used in each case for analysis.

### Analysis to determine mutation effects on cell entry in pseudovirus deep mutational scanning

For the experiments to determine the effects of mutations on cell entry, the Illumina sequencing of the non-integrated lentiviral DNA extracted from the infected cells was analyzed to determine the counts of each barcoded variant in the infected cells in the infections with pseudovirus libraries pseudotyped with the envelope protein variants versus VSV-G. This analysis was performed using dms-vep-pipeline-3 (version 3.27.0, https://github.com/dms-vep/dms-vep-pipeline-3), and all the computer code and barcode counts are in the GitHub repository at https://github.com/dms-vep/CHIKV-181-25-E-DMS with the notebooks performing the analysis rendered at (https://dms-vep.org/CHIKV-181-25-E-DMS/appendix.html.

Briefly, after counting the number of reads corresponding to each barcoded variant in each condition, we filtered only for variants with sufficient counts in the VSV-G only condition to make reliable estimates of the frequencies. We then computed a functional (cell entry) score for each variant as *log_2_([counts of variant in E-pseudotyped libraries / counts of variant in VSV-G-pseudotyped libraries] / [counts of all wildtype variants in E-pseudotyped libraries / counts of all wildtype variants in VSV-G-pseudotyped libraries])*. With this formula, envelope protein variants that have wild-type-like cell entry have scores of zero, and envelope protein variants with impaired cell entry relative to wild-type have scores less than zero. A score of −1 corresponds to two-fold reduced entry, a score of −2 corresponds to four-fold reduced entry, etc; note that we can only reliably measure scores to around −5 or −6 before reaching the lower limit of detection.

After computing the functional (cell entry) scores for all variants, we deconvolved these scores to estimate the effects of individual mutations, since some variants have multiple mutations. To do this, we used a global epistasis model^43^ as implemented in the multidms package^44^, using a sigmoid global epistasis function. The result is an estimate of the effect of each mutation on cell entry; these estimates are highly correlated with the measurements made only using single-mutant variants but have better mutation coverage due to inclusion of the multi-mutant variants (see analysis notebooks in “Functional effects of mutations” section of https://dms-vep.org/CHIKV-181-25-E-DMS/appendix.html). The measurements for each cell type were done with two experimental replicates of each of the two mutant libraries; except for the extended data figures showing the replicate-to-replicate correlations, all plots in this paper show the median measurement across replicates. Filtering is performed to only retain estimates for mutations found in a sufficient number of variants to provide high-confidence measurements. Hovering over the interactive plots at https://dms-vep.org/CHIKV-181-25-E-DMS/cell_entry.html allows you to see the measurements for each individual replicate. For experiments that were done separately with the libraries for each half (E3-E2 and 6K-E1) versus the pool, the analysis was performed separately for each half-library and then the results computationally combined.

### Analysis to determine mutation effects on MXRA8 binding in pseudovirus deep mutational scanning

For the experiments to determine the effects of mutations on MXRA8 binding (as assessed neutralization of pseudovirus by soluble MXRA8-Fc), the Illumina sequencing of the non-integrated lentiviral DNA extract from infected cells was analyzed to determine the counts of each barcoded variant at each MXRA8-Fc concentration alongside the counts of non-neutralized standard VSV-G pseudotyped virus^37^. This analysis was performed using dms-vep-pipeline-3 (version 3.27.0, https://github.com/dms-vep/dms-vep-pipeline-3), and all of computer code and barcode counts are in the GitHub repository at https://github.com/dms-vep/CHIKV-181-25-E-DMS with the notebooks performing the analysis rendered at https://dms-vep.org/CHIKV-181-25-E-DMS/appendix.html.

Briefly, we first converted the barcode counts to the fractional neutralization of each variant at each soluble MXRA8-Fc concentration using the non-neutralized standard counts using a previously described approach^37^. We then analyzed these concentration-dependent neutralization fractions for all variants using a previously described biophysical model^95^ as implemented in the polyclonal software package (version 6.16, https://github.com/jbloomlab/polyclonal), treating the soluble MXRA8-Fc equivalently to an antibody in the framework of the model. The result of analyzing the data with this model is an estimate of the effect of each mutation on MXRA8 binding, and the data are reported such that positive values indicate mutations that increase MXRA8 binding (increase neutralization by soluble MXRA8-Fc). For details on this analysis, see the analysis notebooks in the “MXRA8 binding” section of https://dms-vep.org/CHIKV-181-25-E-DMS/appendix.html. As shown in those notebooks, the measurements were nosier (worse correlations among replicates) for human than mouse MXRA8 binding, so the latter values are more reliable. Hovering over the interactive plots at https://dms-vep.org/CHIKV-181-25-E-DMS/MXRA8_binding.html allows you to see the measurements for each individual replicate.

### Structural visualizations

UCSF ChimeraX v1.10.1^96^ was used for structural visualizations in the paper figures, using the Protein Data Bank accession IDs given in the figure legends. The interactive structural visualizations at https://dms-vep.org/CHIKV-181-25-E-DMS/structures.html were created using dms-viz^97^.

## Extended Data Figures

**Extended Data Figure 1.**
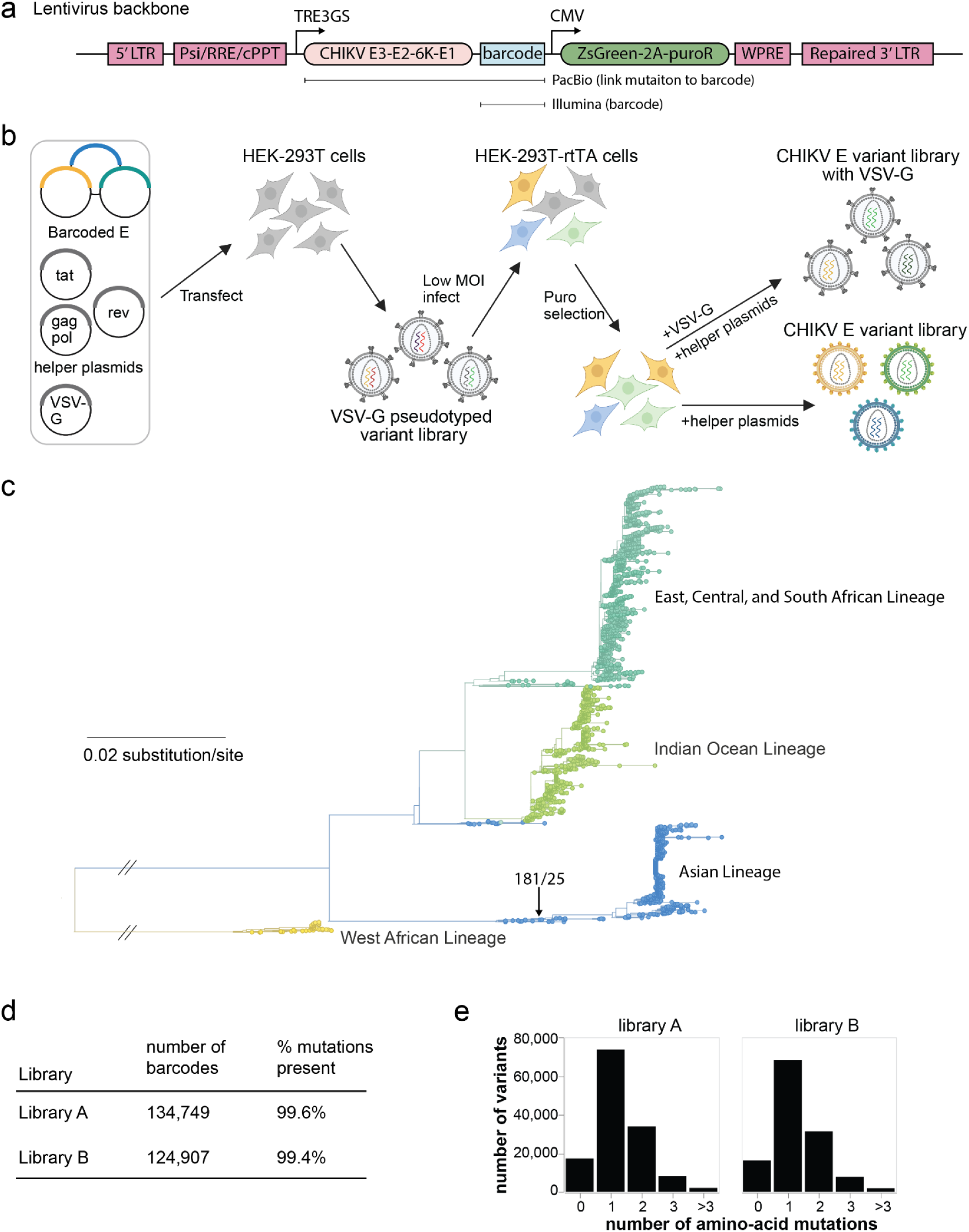
Pseudovirus deep mutational scanning of CHIKV envelope proteins. **(a)** Schematic of the lentiviral genome (backbone) used for the pseudovirus deep mutational scanning^37^. The backbone contains standard elements (the 5’ LTR, packing signal Psi, RRE, cPPT, 3’ LTR) with the 3’ LTR full-length (not deleted) to allow reactivation of integrated proviruses. The backbone also encodes the CHIKV envelope proteins (E3-E2-6K-E1) under an inducible TRE3GS promoter followed by a random nucleotide barcode after the stop codon. A constitutive CMV promoter drives expression of ZsGreen and a puromycin resistance gene. **(b)** Two-step strategy to produce genotype-phenotype linked library of pseudotyped lentiviral particles^37^. 293T cells are transfected with a library of backbone plasmids as well helper plasmids encoding Tat, Gag-Pol, and Rev, and a plasmid encoding VSV-G. The resulting virions are used to infect 293T cells expressing the rtTA protein needed for the inducible TRE3GS promoter at low multiplicity of infection, and cells with integrants are selected using puromycin. These cells now store the lentiviral genome library, and genotype-phenotype linked pseudoviruses expressing the CHIKV envelope proteins can be generated by transfecting the helper plasmids. To determine the library composition independent of the function of the CHIKV envelope proteins, we also generate control virions expressing VSV-G. **(c)** Phylogenetic tree of genes encoding the CHIKV structural proteins. The 181/25 strain used for our libraries is labeled, and the tree is colored by the four major clades. The tree was built on an alignment of the nucleotide sequences for the structural proteins (Capsid, E3, E2, 6K, E1), as downloaded from NCBI virus on April 4th, 2025. See https://dms-vep.org/CHIKV-181-25-E-DMS/tree.html for an interactive version of this phylogenetic tree. **(d)** Number of uniquely barcoded E protein variants in each of the two replicate libraries, and the percentage of all possible amino-acid mutations that they cover. **(e)** Number of amino-acid mutations per variant in the CHIKV envelope proteins across all barcoded variants in each library.

**Extended Data Figure 2.**
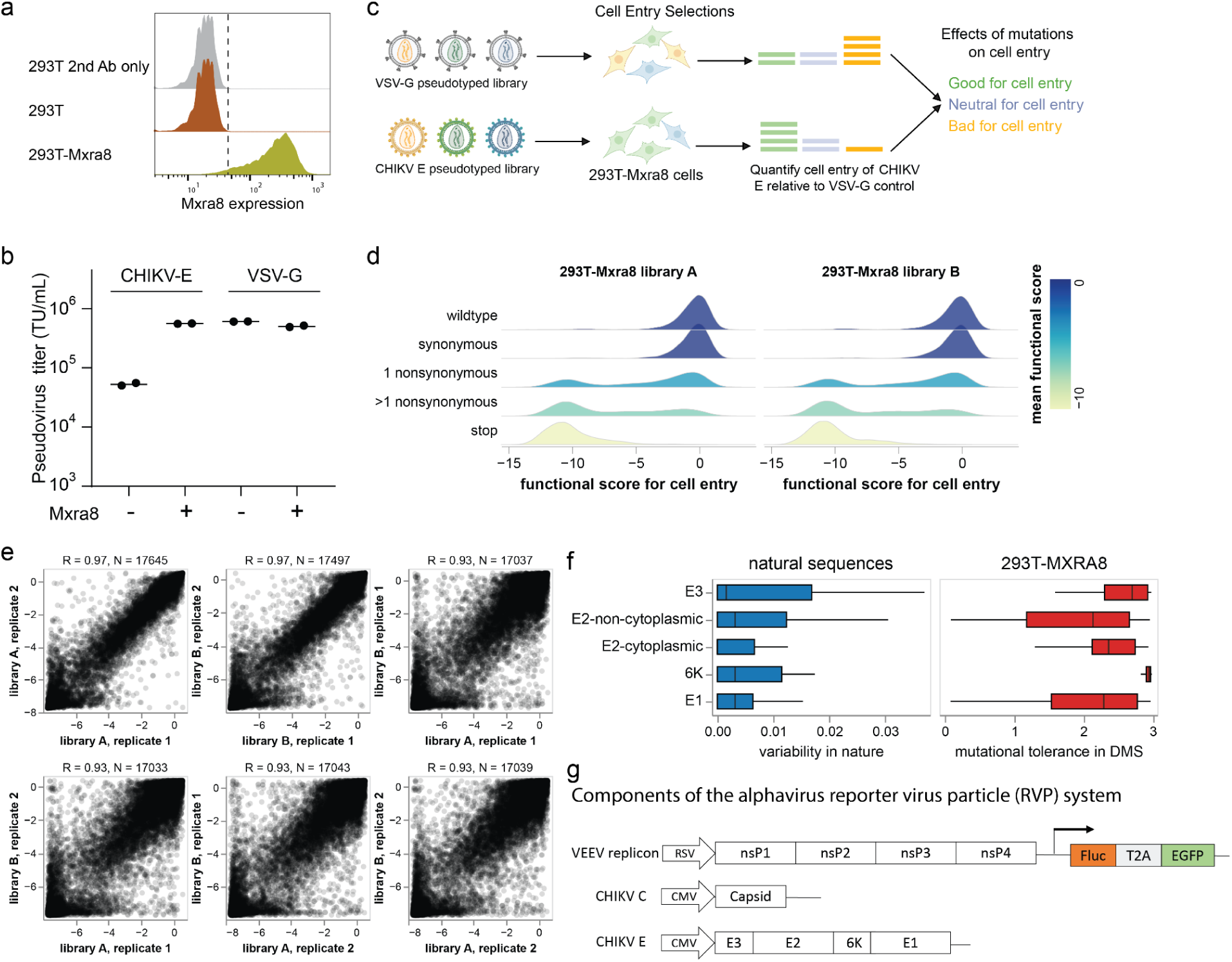
Experiments to measure effects of mutations on entry in 293T-MXRA8 cells. **(a)** Immunostaining of MXRA8 expression in 293T-MXRA8 cells. Flow cytometry analysis of 293T cells stained with fluorophore-conjugated secondary antibody only (gray), 293T cells stained with anti-MXRA8 primary antibody and secondary antibody (brown) and 293T-MXRA8 cells stained with anti-MXRA8 primary antibody and secondary antibody (yellow). **(b)** Titers of pseudoviruses (lentiviral pseudotyped particles) expressing the CHIKV envelope proteins from the 181/25 strain or VSV-G on 293T or 293T-MXRA8 target cells, quantified as transducing units (TU) per ml. **(c)** Overview of deep mutational scanning for entry into 293T-MXRA8 cells. Cells are infected with VSV-G pseudotyped or envelope proteins-pseudotyped variant library, produced as shown in **Extended Data Fig. 1b**. At 12 hours post infection, unintegrated viral DNA is extracted from the infected cells^37,99^ and variant barcodes are sequenced. All variants infect cells when VSV-G is present, but only variants with functional envelope proteins infect cells when VSV-G is absent. A cell entry score for each variant is calculated by comparing the barcode frequency between CHIKV envelope proteins pseudotyped library and VSV-G pseudotyped library. **(d)** Distributions of cell-entry scores for variants with different types of mutations. Values of zero indicate wildtype-like cell entry and values less than zero indicate impaired cell entry. **(e)** Correlations of per-mutation effects on cell entry measured for the two technical replicates of each of the two independent libraries after deconvolution of variant functional scores using global epistasis models (see Methods). Technical replicates of the same library are very well correlated (R = 0.97), and replicates of the independent libraries are also well correlated (R = 0.93). **(f)** Variability of sites in different regions of the envelope proteins among natural sequences as quantified by the per-site Shannon entropy across an alignment of natural CHIKV sequences versus mutational tolerance in pseudovirus deep mutational scanning quantified as the entropy of the measured amino-acid preferences for each site. Higher values indicate more variability or mutational tolerance. **(g)** Schematic of single-cycle alphavirus reporter virus particle (RVP) system. One plasmid encodes a replicon of the attenuated VEEV TC-83 strain with the viral structural proteins replaced by firefly luciferase (Fluc) and enhanced green fluorescent protein (EGFP). The other two plasmids encode the CHIKV capsid and envelope proteins separately. Co-transfection of these three plasmids into cells produces RVP that can only infect target cells once since they do not encode any of the structural proteins.

**Extended Data Figure 3.**
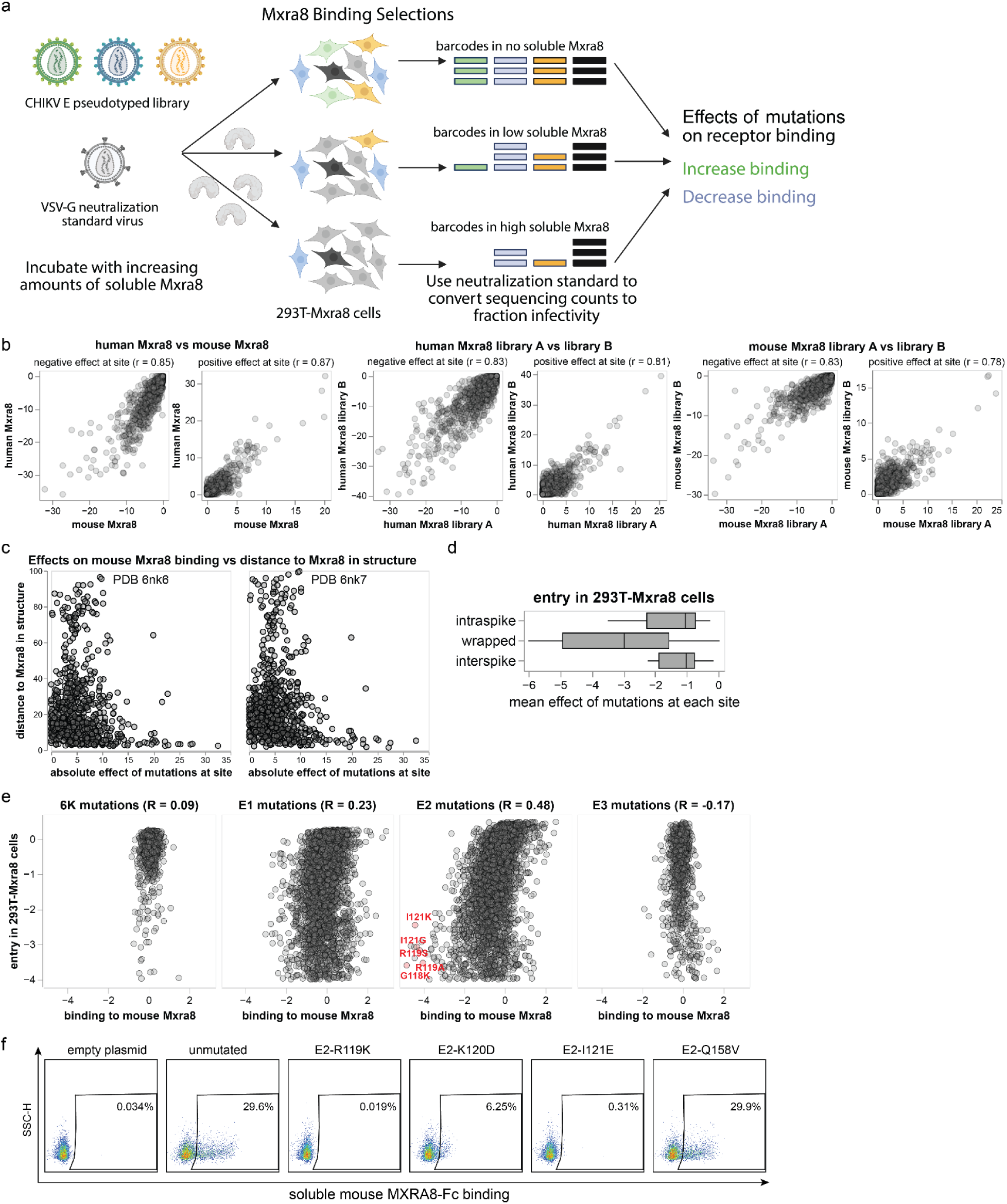
Experiments to measure effects of mutations on MXRA8 binding. **(a)** Deep mutational scanning to measure MXRA8 binding. Pseudovirus libraries expressing the CHIKV envelope protein mutants are mixed with a neutralization ‘‘standard’’ pseudovirus^37^ that expresses only VSV-G (and so is not affected by MXRA8) and contains defined barcodes in its genome. Libraries mixed with the neutralization standard are used to infect 293T-MXRA8 in the presence of no soluble MXRA8-Fc or increasing concentrations of soluble MXRA8-Fc, which neutralizes infection by the CHIKV envelope protein expressing pseudoviruses. Barcodes of infecting viruses are sequenced, and the sequencing counts are converted to fractional infectivity of each pseudovirus mutant at each MXRA8-Fc concentration using the neutralization standard^37^. The data are analyzed to determine the effect of each mutation on binding (see **Methods**). **(b)** Correlations of summed effects of all mutations at each site on binding to human MXRA8 and mouse MXRA8 binding (taking the mean effect of mutations across the two deep mutational scanning libraries) as well as correlations between the effects of mutations on binding as measured in each of the two libraries for the same MXRA8. The sums and correlations are computed separately for mutations that decrease or increase MXRA8 binding. **(c)** Sum of absolute value of mutation effects at each site on mouse MXRA8 binding at each site versus the distance of that site to mouse MXRA8 in two cryo-EM structures (PDB identifiers 6nk6 and 6nk7)^51^. **(d)** Mean effect on cell entry in 293T-MXRA8 cells of mutations at sites that are involved in binding to MXRA8 via different modes. **(e)** Correlation of effects of mutations on entry into 293T-MXRA8 cells and binding to mouse MXRA8, computed separately for each envelope protein. Key E2 mutations that strongly impair MXRA8 binding and entry into 293T-MXRA8 cells are labeled and highlighted in red. R indicates the Pearson correlation. **(f)** Representative flow cytometry plots of the cell-based binding assay showing mouse MXRA8-Fc binding to cells at 1.56 μg/mL; this plot represents one of the concentration points on the curve shown in Figure 3e.

**Extended Data Figure 4.**
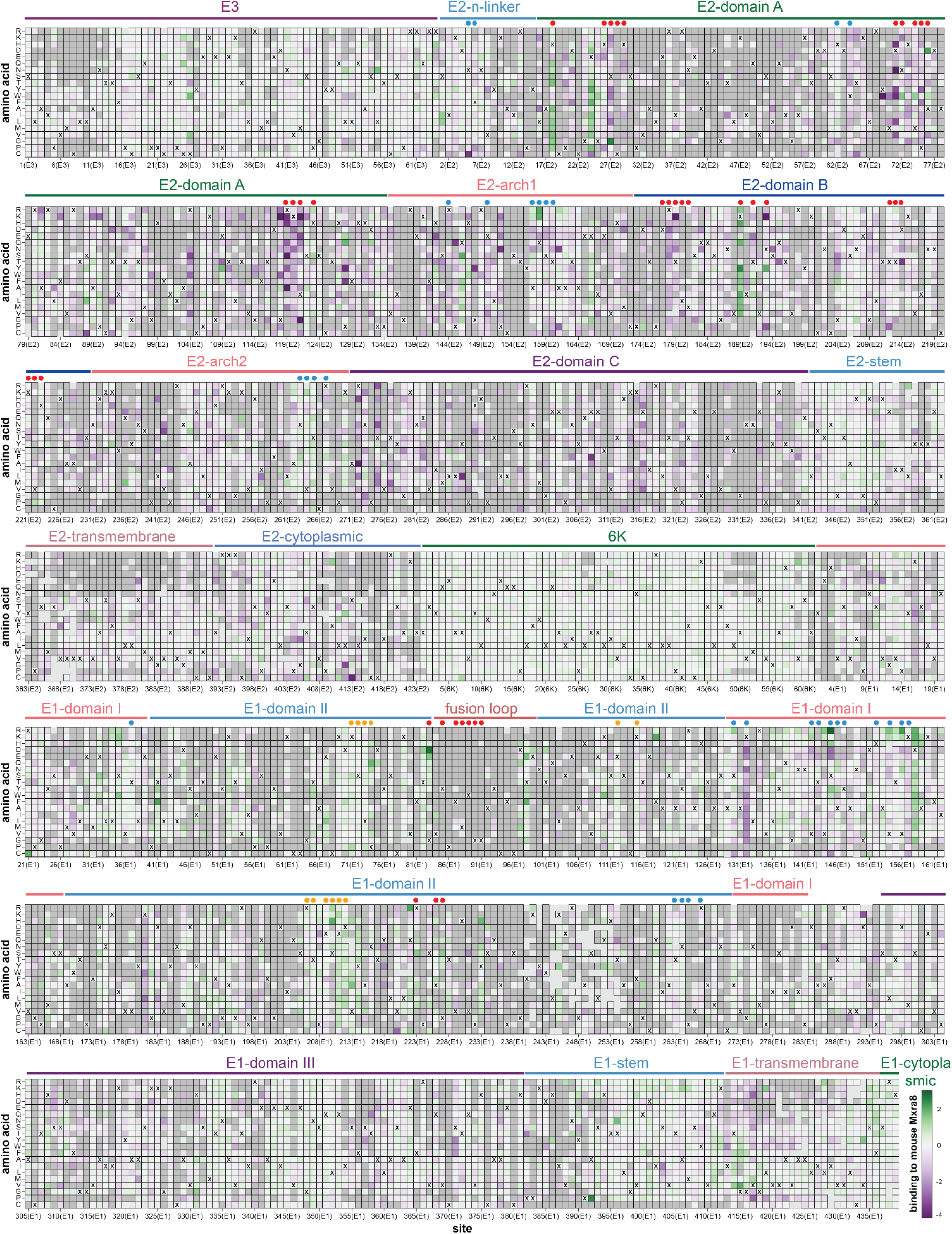
Effects of mutations on binding to mouse MXRA8. Mutations that are colored purple reduce MXRA8 binding, while mutations colored green improve MXRA8 binding. Dark gray squares indicate mutations that ar e too deleterious for cell entry (effect < −4) to measure their effect on MXRA8 binding. Light grey squares indicate mutations with missing measurements. MXRA8-contact sites as annotated by Basore et al^51^ are indicated with a dot above the heat map (red dot for wrapped contact sites, blue dot for intraspike contact sites, and orange dot for interspike contact sites). The amino-acid identity in the parental 181/25 strain at each site is represented with an ‘‘X’’. See https://dms-vep.org/CHIKV-181-25-E-DMS/MXRA8_binding.html for an interactive version of this heatmap that is easier to explore than this static version; that link also provides a heatmap of how mutations affect binding to human MXRA8.

**Extended Data Figure 5.**
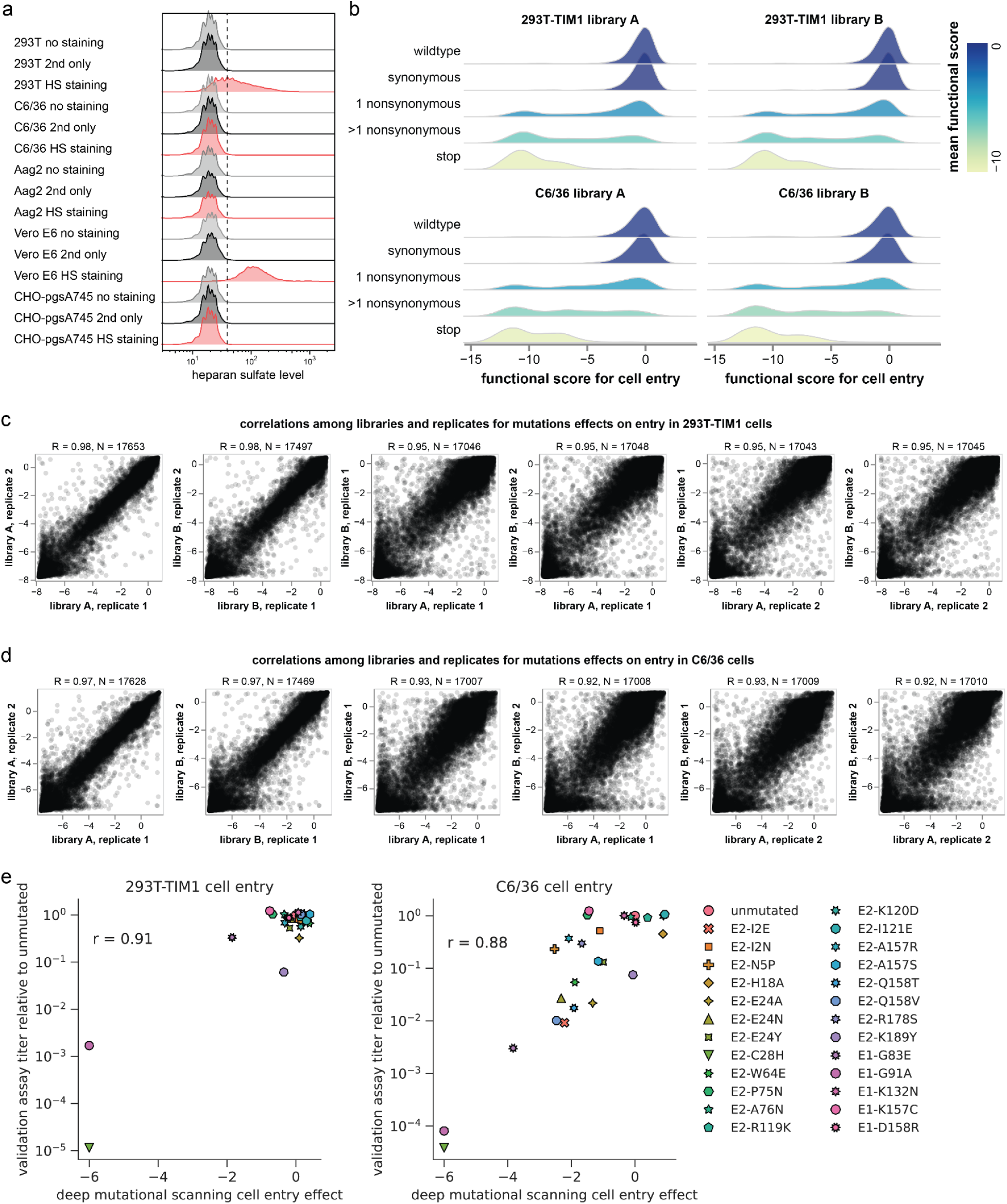
Experiments to measure effects of mutations on entry in 293T-TIM1 and C6/36 cells. (**a**) Immunostaining of heparan sulfate expression in different cell lines. Flow cytometry analysis of cells without staining (grey), cells stained with fluorophore-conjugated secondary antibody only (black), and cells stained with anti-heparan sulfate primary antibody and secondary antibody (red). Vero E6 cells express high levels of heparan sulfate and CHO-pgsA745 cells do not express heparan sulfate, and so are included as controls^24,77^. (**b**) Effects of different types of mutations on cell entry; these data are comparable to those shown in **Extended Data Fig. 2d** except for 293T-TIM1 and C6/36 cells. (**c**), (**d**) Correlations among effects of mutations on cell entry measured in different deep mutational scanning replicates; these data are comparable to those shown in **Extended Data Fig. 2e** except for 293T-TIM1 or C6/36 cells. (**e**) Validation of deep mutational scanning measurements of mutational effects on cell entry using alphavirus RVPs; these data are comparable to those shown in Fig. 2d except for 293T-TIM1 and C6/36 cells.

**Extended Data Figure 6.**
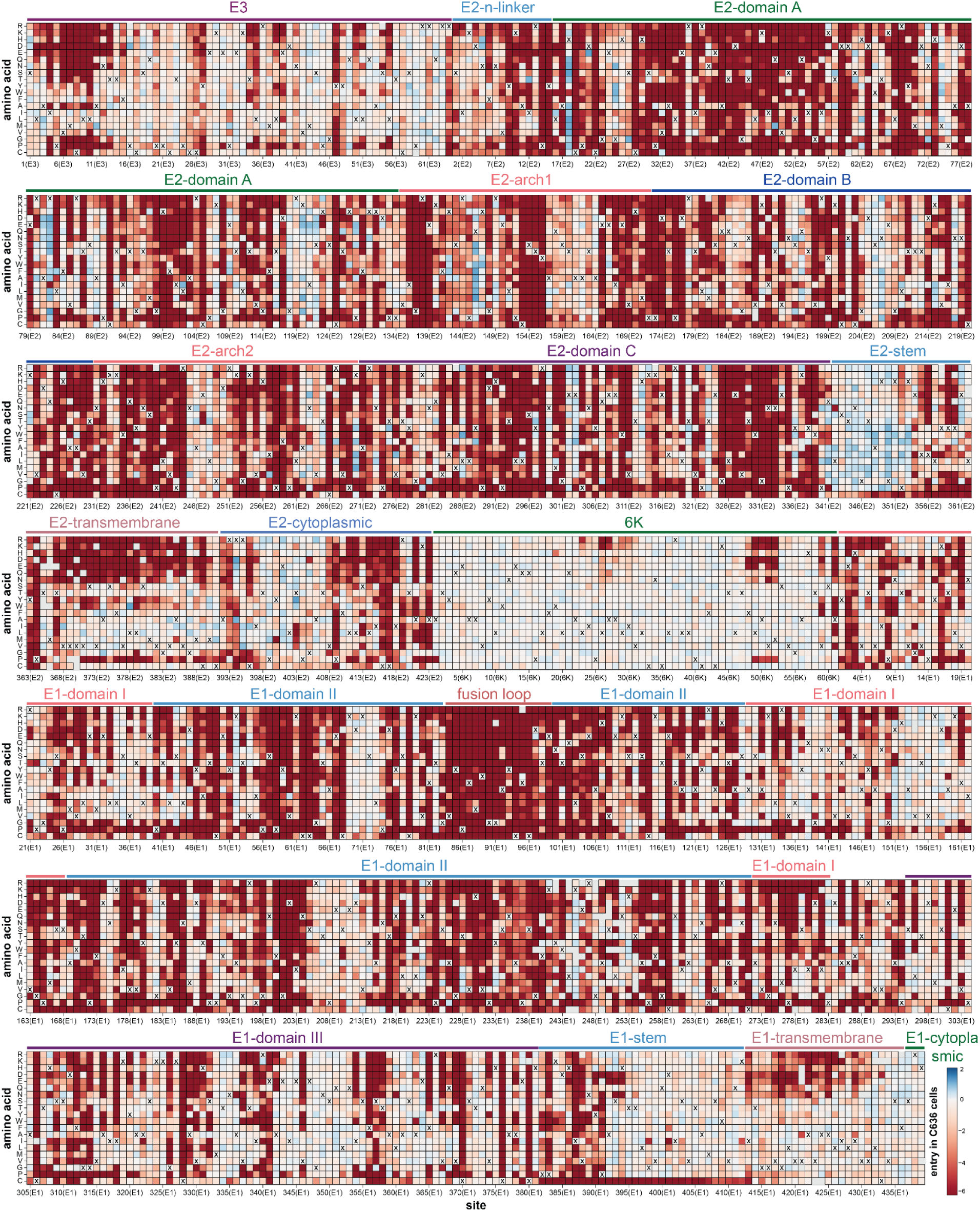
Effects of mutations on entry in C6/36 cells. Comparable to Fig. 1 except for entry in C6/36 cells instead of 293T-MXRA8 cells. See https://dms-vep.org/CHIKV-181-25-E-DMS/cell_entry.html for an interactive version.

**Extended Data Figure 7.**
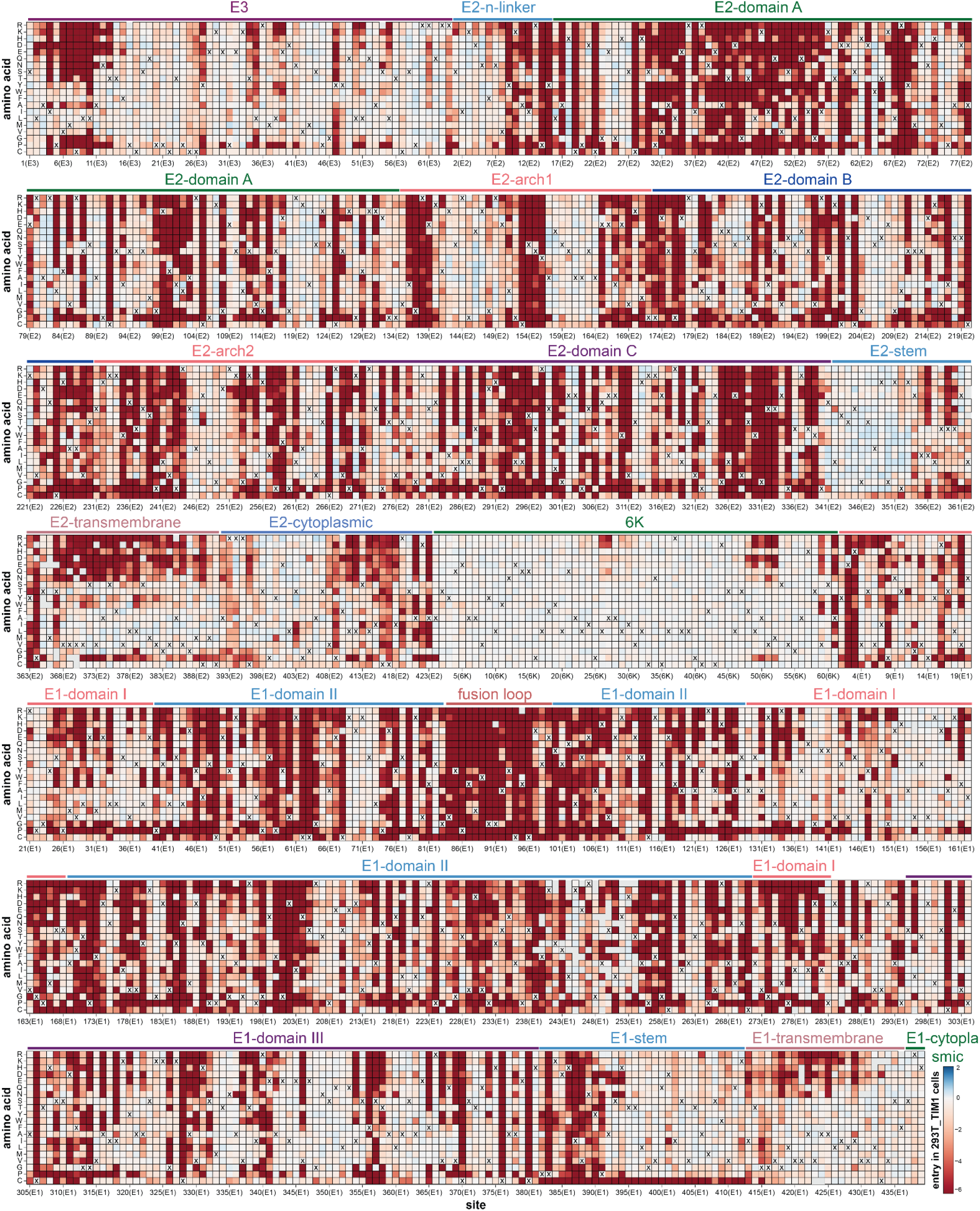
Effects of mutations on entry into 293T-TIM1 cells. Comparable to Fig. 1 except for entry in 293T-TIM1 cells instead of 293T-MXRA8 cells. See https://dms-vep.org/CHIKV-181-25-E-DMS/cell_entry.html for an interactive version.

**Extended Data Figure 8.**
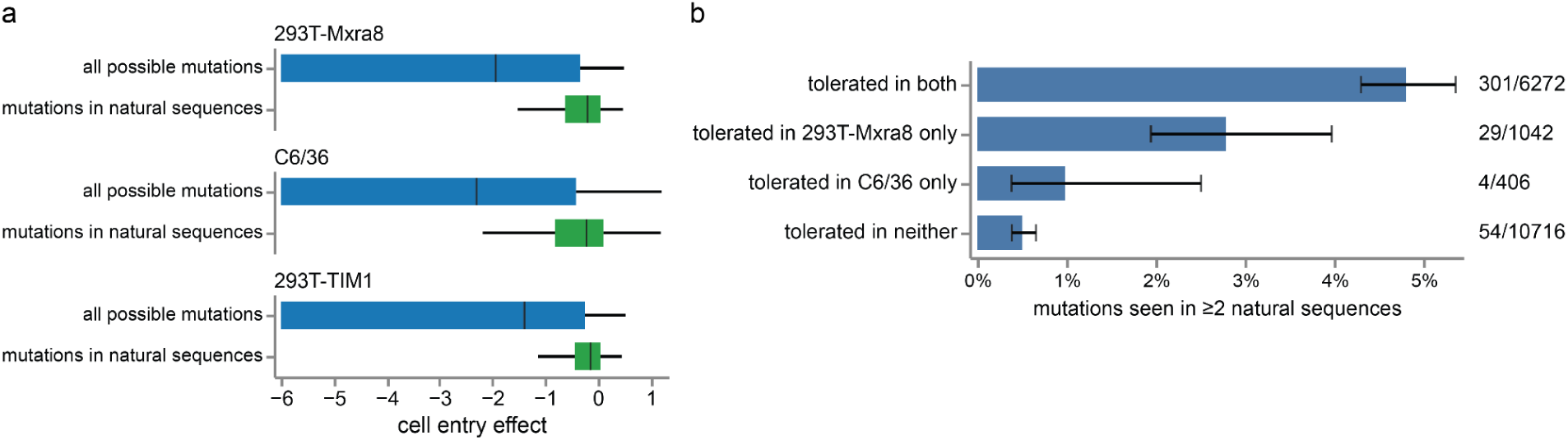
Measured effects on cell entry of mutations in natural CHIKV sequences. (a) Effects on pseudovirus entry in each cell line as measured in the deep mutational scanning of all possible single amino-acid mutations to the envelope proteins and just the mutations observed in natural CHIKV sequences. (b) Percent of all possible mutations that are observed in at least two natural CHIKV sequences, stratified by if the mutation is tolerated in one or both cell lines. A mutation is classified as tolerated if its effect on entry into that cell line as measured in the deep mutational scanning is ≥ −1. The numbers to the right of each bar gives the number of mutations in that class observed in natural sequences over the total number in the class (e.g., 301/6272 means that 301 of the 6,272 mutations that are tolerated in both cell lines are observed at least twice among natural CHIKV sequences).

**Extended Data Figure 9.**
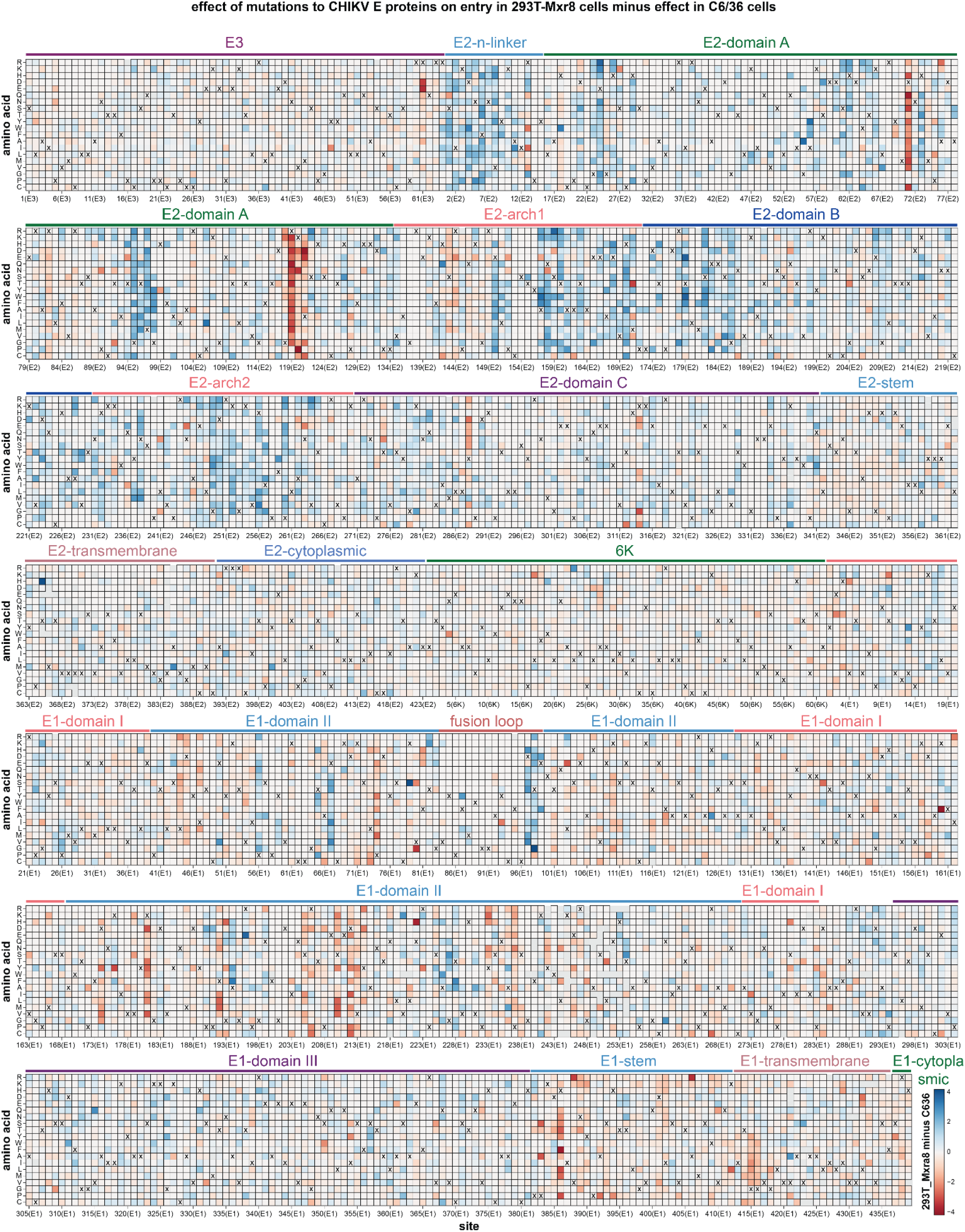
Differences in effects of mutations on entry in 293T-MXRA8 versus C6/36 cells. Each square in the heatmap represents the effect of the mutation on entry in 293T-MXRA8 cells minus the effect on entry in C6/36 cells. Positive values represent mutations that are more favorable for entry in 293T-MXRA8 than C6/36 cells, values of zero indicate mutations with similar effects on entry in both cells, and negative values represent mutations that are more favorable for entry in C6/36 versus 293T-MXRA8 cells. The wildtype amino acid in the 181/25 strain at each site is indicated with an “X”. The handful of gray squares indicate mutations that were not measured with high confidence in the deep mutational scanning. See https://dms-vep.org/CHIKV-181-25-E-DMS/cell_entry_diffs.html for interactive visualization of the same data.

**Extended Data Figure 10.**
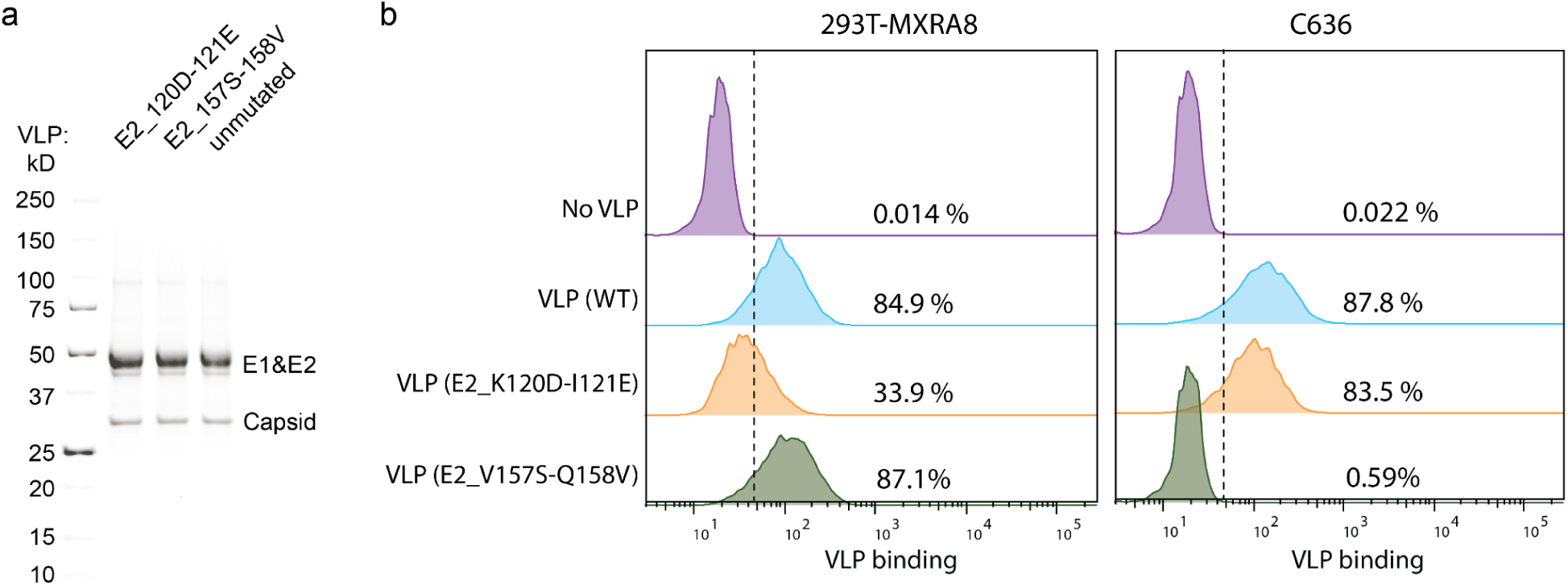
Cell-based binding assay to measure the interaction between CHIKV VLPs and cells. (**a**) SDS-PAGE analysis of the purified CHIKV VLPs from the 37997 strain under reducing conditions. (**b**) Cell-based binding assay to detect the interaction between cells and VLPs. Representative flow cytometry plots are shown. The VLPs are incubated with the cells, and then VLP binding to each is quantified using a mixture of antibodies specific to the CHIKV envelope proteins (see Methods for details). Note that the 37997 E2 has a V at site 157 whereas the 181/25 E2 has an A at site 157, which is why the mutation is V157S here and A157S elsewhere in the paper.

## Supplementary Files

Supplementary File 1. Effects of mutations on entry in each cell and on Mxra8 binding.

This supplementary file CSV is also available at https://github.com/dms-vep/CHIKV-181-25-E-DMS/blob/main/results/summaries/entry_293T-Mxra8_C636_293T-TIM1_Mxra8-binding.csv

## References

1. Chopra, A., Anuradha, V., Ghorpade, R. & Saluja, M. Acute Chikungunya and persistent musculoskeletal pain following the 2006 Indian epidemic: a 2-year prospective rural community study. Epidemiol Infect 140, 842–850 (2012).

2. Schilte, C. et al. Chikungunya virus-associated long-term arthralgia: a 36-month prospective longitudinal study. PLoS Negl Trop Dis 7, e2137 (2013).

3. Schwartz, O. & Albert, M. L. Biology and pathogenesis of chikungunya virus. Nat Rev Microbiol 8, 491–500 (2010).

4. Silva, L. A. & Dermody, T. S. Chikungunya virus: epidemiology, replication, disease mechanisms, and prospective intervention strategies. J Clin Invest 127, 737–749 (2017).

5. Tsetsarkin, K. A., Vanlandingham, D. L., McGee, C. E. & Higgs, S. A single mutation in chikungunya virus affects vector specificity and epidemic potential. PLoS Pathog 3, e201 (2007).

6. Vazeille, M. et al. Two Chikungunya isolates from the outbreak of La Reunion (Indian Ocean) exhibit different patterns of infection in the mosquito, Aedes albopictus. PLoS One 2, e1168 (2007).

7. Tsetsarkin, K. A. & Weaver, S. C. Sequential adaptive mutations enhance efficient vector switching by Chikungunya virus and its epidemic emergence. PLoS Pathog 7, e1002412 (2011).

8. Farooq, Z. et al. Impact of climate and Aedes albopictus establishment on dengue and chikungunya outbreaks in Europe: a time-to-event analysis. Lancet Planet Health 9, e374–e383 (2025).

9. Dai, Z. et al. Global Assessment of current and future chikungunya virus transmission risk using optimized maxent modeling. Acta Trop 269, 107756 (2025).

10. de Souza, W. M., Lecuit, M. & Weaver, S. C. Chikungunya virus and other emerging arthritogenic alphaviruses. Nat. Rev. Microbiol. https://doi.org/10.1038/s41579-025-01177-8 (2025) doi:10.1038/s41579-025-01177-8.

11. Yap, M. L. et al. Structural studies of Chikungunya virus maturation. Proc Natl Acad Sci U S A 114, 13703–13707 (2017).

12. Voss, J. E. et al. Glycoprotein organization of Chikungunya virus particles revealed by X-ray crystallography. Nature 468, 709–712 (2010).

13. Kim, A. S. & Diamond, M. S. A molecular understanding of alphavirus entry and antibody protection. Nat Rev Microbiol 21, 396–407 (2023).

14. Byrd, E. A. & Kielian, M. The Alphavirus E2 Membrane-Proximal Domain Impacts Capsid Interaction and Glycoprotein Lattice Formation. J Virol 93, (2019).

15. Zhang, R. et al. Mxra8 is a receptor for multiple arthritogenic alphaviruses. Nature 557, 570–574 (2018).

16. Zhang, R. et al. Expression of the Mxra8 Receptor Promotes Alphavirus Infection and Pathogenesis in Mice and Drosophila. Cell Rep 28, 2647–2658.e5 (2019).

17. Yin, P. et al. Chikungunya virus cell-to-cell transmission is mediated by intercellular extensions in vitro and in vivo. Nat Microbiol 8, 1653–1667 (2023).

18. Reyes Ballista, J. M., et al. Chikungunya virus entry and infectivity is primarily facilitated through cell line dependent attachment factors in mammalian and mosquito cells. Front Cell Dev Biol 11, 1085913 (2023).

19. Fongsaran, C. et al. Involvement of ATP synthase β subunit in chikungunya virus entry into insect cells. Arch Virol 159, 3353–3364 (2014).

20. Ghosh, A., Desai, A., Ravi, V., Narayanappa, G. & Tyagi, B. K. Chikungunya Virus Interacts with Heat Shock Cognate 70 Protein to Facilitate Its Entry into Mosquito Cell Line. Intervirology 60, 247–262 (2017).

21. Kirui, J. et al. The Phosphatidylserine Receptor TIM-1 Enhances Authentic Chikungunya Virus Cell Entry. Cells 10, (2021).

22. Silva, L. A. et al. A single-amino-acid polymorphism in Chikungunya virus E2 glycoprotein influences glycosaminoglycan utilization. J Virol 88, 2385–2397 (2014).

23. Tanaka, A. et al. Genome-Wide Screening Uncovers the Significance of N-Sulfation of Heparan Sulfate as a Host Cell Factor for Chikungunya Virus Infection. J Virol 91, (2017).

24. McAllister, N. et al. Chikungunya Virus Strains from Each Genetic Clade Bind Sulfated Glycosaminoglycans as Attachment Factors. J Virol 94, (2020).

25. Kose, N. et al. A lipid-encapsulated mRNA encoding a potently neutralizing human monoclonal antibody protects against chikungunya infection. Sci Immunol 4, (2019).

26. Fox, J. M. et al. Broadly Neutralizing Alphavirus Antibodies Bind an Epitope on E2 and Inhibit Entry and Egress. Cell 163, 1095–1107 (2015).

27. Quiroz, J. A. et al. Human monoclonal antibodies against chikungunya virus target multiple distinct epitopes in the E1 and E2 glycoproteins. PLoS Pathog 15, e1008061 (2019).

28. Raju, S. et al. A chikungunya virus-like particle vaccine induces broadly neutralizing and protective antibodies against alphaviruses in humans. Sci Transl Med 15, eade8273 (2023).

29. Ribeiro Dos Santos, G., et al. Global burden of chikungunya virus infections and the potential benefit of vaccination campaigns. Nat Med 31, 2342–2349 (2025).

30. Weber, W. C., Streblow, D. N. & Coffey, L. L. Chikungunya Virus Vaccines: A Review of IXCHIQ and PXVX0317 from Pre-Clinical Evaluation to Licensure. BioDrugs 38, 727–742 (2024).

31. Zhou, Q. F. et al. Structural basis of Chikungunya virus inhibition by monoclonal antibodies. Proc Natl Acad Sci U S A 117, 27637–27645 (2020).

32. Smith, S. A. et al. Isolation and Characterization of Broad and Ultrapotent Human Monoclonal Antibodies with Therapeutic Activity against Chikungunya Virus. Cell Host Microbe 18, 86–95 (2015).

33. Zumla, A., Ntoumi, F., Ippolito, G. & PANDORA-ID-NET Consortium. Chikungunya virus disease returns to Europe: a turning point for the global arboviral landscape. Lancet https://doi.org/10.1016/S0140-6736(25)01458-8 (2025) doi:10.1016/S0140-6736(25)01458-8.

34. Bartholomeeusen, K. et al. Chikungunya fever. Nat Rev Dis Primers 9, 17 (2023).

35. Kuo, L. & Dong, J. With Drones and ‘Elephant Mosquitoes,’ China Wages All-Out War on a Virus. The New York Times (2025).

36. AFP. Outbreak of Chikungunya Virus Poses Global Risk, Warns WHO. ScienceAlert https://www.sciencealert.com/outbreak-of-chikungunya-virus-poses-global-risk-warns-who (2025).

37. Dadonaite, B. et al. A pseudovirus system enables deep mutational scanning of the full SARS-CoV-2 spike. Cell 186, 1263–1278.e20 (2023).

38. Larsen, B. B. et al. Functional and antigenic landscape of the Nipah virus receptor-binding protein. Cell 188, 2480–2494.e22 (2025).

39. Levitt, N. H. et al. Development of an attenuated strain of chikungunya virus for use in vaccine production. Vaccine 4, 157–162 (1986).

40. Gorchakov, R. et al. Attenuation of Chikungunya virus vaccine strain 181/clone 25 is determined by two amino acid substitutions in the E2 envelope glycoprotein. J Virol 86, 6084–6096 (2012).

41. Snyder, J. E. et al. Functional characterization of the alphavirus TF protein. J Virol 87, 8511–8523 (2013).

42. Kim, A. S. et al. An Evolutionary Insertion in the Mxra8 Receptor-Binding Site Confers Resistance to Alphavirus Infection and Pathogenesis. Cell Host Microbe 27, 428–440.e9 (2020).

43. Otwinowski, J., McCandlish, D. M. & Plotkin, J. B. Inferring the shape of global epistasis. Proc Natl Acad Sci U S A 115, E7550–E7558 (2018).

44. Haddox, H. K., et al. Jointly modeling deep mutational scans identifies shifted mutational effects among SARS-CoV-2 spike homologs. bioRxiv (2023) doi:10.1101/2023.07.31.551037.

45. Li, L., Jose, J., Xiang, Y., Kuhn, R. J. & Rossmann, M. G. Structural changes of envelope proteins during alphavirus fusion. Nature 468, 705–708 (2010).

46. Uchime, O., Fields, W. & Kielian, M. The role of E3 in pH protection during alphavirus assembly and exit. J Virol 87, 10255–10262 (2013).

47. Melton, J. V. et al. Alphavirus 6K proteins form ion channels. J Biol Chem 277, 46923–46931 (2002).

48. Sanz, M. A., Madan, V., Carrasco, L. & Nieva, J. L. Interfacial domains in Sindbis virus 6K protein. Detection and functional characterization. J Biol Chem 278, 2051–2057 (2003).

49. Ren, S. C., Qazi, S. A., Towell, B., Wang, J. C.-Y. & Mukhopadhyay, S. Mutations at the Alphavirus E1’-E2 Interdimer Interface Have Host-Specific Phenotypes. J Virol 96, e0214921 (2022).

50. Zhang, R. et al. 4.4 Å cryo-EM structure of an enveloped alphavirus Venezuelan equine encephalitis virus. EMBO J 30, 3854–3863 (2011).

51. Basore, K. et al. Cryo-EM Structure of Chikungunya Virus in Complex with the Mxra8 Receptor. Cell 177, 1725–1737.e16 (2019).

52. Song, H. et al. Molecular Basis of Arthritogenic Alphavirus Receptor MXRA8 Binding to Chikungunya Virus Envelope Protein. Cell 177, 1714–1724.e12 (2019).

53. Clark, L. E. et al. VLDLR and ApoER2 are receptors for multiple alphaviruses. Nature 602, 475–480 (2022).

54. Li, W. et al. Shifts in receptors during submergence of an encephalitic arbovirus. Nature 632, 614–621 (2024).

55. Wang, Q. et al. Antigenic characterization of the SARS-CoV-2 Omicron subvariant BA.2.75. Cell Host Microbe 30, 1512–1517.e4 (2022).

56. Cao, Y. et al. Imprinted SARS-CoV-2 humoral immunity induces convergent Omicron RBD evolution. Nature 614, 521–529 (2023).

57. Dadonaite, B. et al. Spike deep mutational scanning helps predict success of SARS-CoV-2 clades. Nature 631, 617–626 (2024).

58. Igarashi, A. Isolation of a Singh’s Aedes albopictus cell clone sensitive to Dengue and Chikungunya viruses. J Gen Virol 40, 531–544 (1978).

59. Roberts, G. C. et al. Evaluation of a range of mammalian and mosquito cell lines for use in Chikungunya virus research. Sci Rep 7, 14641 (2017).

60. Mohamed Ali, S., Amroun, A., de Lamballerie, X. & Nougairède, A. Evolution of Chikungunya virus in mosquito cells. Sci Rep 8, 16175 (2018).

61. Lee, R. C. H. & Chu, J. J. H. Proteomics profiling of chikungunya-infected Aedes albopictus C6/36 cells reveal important mosquito cell factors in virus replication. PLoS Negl. Trop. Dis. 9, e0003544 (2015).

62. Novelo, M. et al. Dengue and chikungunya virus loads in the mosquito Aedes aegypti are determined by distinct genetic architectures. PLoS Pathog. 19, e1011307 (2023).

63. Qu, J. et al. The Hsf1-sHsp cascade has pan-antiviral activity in mosquito cells. *Commun*. Biol. 8, 123 (2025).

64. Simonich, C. A. L. et al. RSV F evolution escapes some monoclonal antibodies but does not strongly erode neutralization by human polyclonal sera. J Virol e0053125 (2025).

65. Jemielity, S. et al. TIM-family proteins promote infection of multiple enveloped viruses through virion-associated phosphatidylserine. PLoS Pathog 9, e1003232 (2013).

66. Moller-Tank, S., Kondratowicz, A. S., Davey, R. A., Rennert, P. D. & Maury, W. Role of the phosphatidylserine receptor TIM-1 in enveloped-virus entry. J Virol 87, 8327–8341 (2013).

67. Holmes, A. C., Basore, K., Fremont, D. H. & Diamond, M. S. A molecular understanding of alphavirus entry. PLoS Pathog 16, e1008876 (2020).

68. Zimmerman, O., Holmes, A. C., Kafai, N. M., Adams, L. J. & Diamond, M. S. Entry receptors - the gateway to alphavirus infection. J Clin Invest 133, (2023).

69. Lan, Q. & Fallon, A. M. Small heat shock proteins distinguish between two mosquito species and confirm identity of their cell lines. Am. J. Trop. Med. Hyg. 43, 669–676 (1990).

70. Fredericks, A. C. et al. Aedes aegypti (Aag2)-derived clonal mosquito cell lines reveal the effects of pre-existing persistent infection with the insect-specific bunyavirus Phasi Charoen-like virus on arbovirus replication. PLoS Negl. Trop. Dis. 13, e0007346 (2019).

71. Akahata, W. & Nabel, G. J. A specific domain of the Chikungunya virus E2 protein regulates particle formation in human cells: implications for alphavirus vaccine design. J Virol 86, 8879–8883 (2012).

72. Stumpf, M. M., Brunetti, T., Davenport, B. J., McCarthy, M. K. & Morrison, T. E. Deep mutationally scanned CHIKV E3/E2 virus library maps viral amino acid preferences and predicts viral escape mutants of neutralizing CHIKV antibodies. J Virol 99, e0008125 (2025).

73. Zimmerman, O. et al. Vertebrate-class-specific binding modes of the alphavirus receptor MXRA8. Cell 186, 4818–4833.e25 (2023).

74. Ma, H. et al. LDLRAD3 is a receptor for Venezuelan equine encephalitis virus. Nature 588, 308–314 (2020).

75. Palakurty, S. et al. The VLDLR entry receptor is required for the pathogenesis of multiple encephalitic alphaviruses. Cell Rep 43, 114809 (2024).

76. Zhai, X. et al. LDLR is used as a cell entry receptor by multiple alphaviruses. Nat Commun 15, 622 (2024).

77. Ashbrook, A. W. et al. Residue 82 of the Chikungunya virus E2 attachment protein modulates viral dissemination and arthritis in mice. J. Virol. 88, 12180–12192 (2014).

78. Negrete, O. A. et al. EphrinB2 is the entry receptor for Nipah virus, an emergent deadly paramyxovirus. Nature 436, 401–405 (2005).

79. Li, W. et al. Angiotensin-converting enzyme 2 is a functional receptor for the SARS coronavirus. Nature 426, 450–454 (2003).

80. Plung, J. S., et al. Lachesin is a mosquito receptor for multiple arthritogenic alphaviruses. bioRxiv (2026) doi:10.64898/2026.07.28.741058.

81. Fan, X., et al. Molecular basis for chikungunya virus recognition of a mosquito-specific receptor. bioRxiv (2026) doi:10.64898/2026.07.28.741352.

82. Edelman, R. et al. Phase II safety and immunogenicity study of live chikungunya virus vaccine TSI-GSD-218. Am J Trop Med Hyg 62, 681–685 (2000).

83. Haddox, H. K., Dingens, A. S., Hilton, S. K., Overbaugh, J. & Bloom, J. D. Mapping mutational effects along the evolutionary landscape of HIV envelope. Elife 7, (2018).

84. Starr, T. N. et al. Shifting mutational constraints in the SARS-CoV-2 receptor-binding domain during viral evolution. Science 377, 420–424 (2022).

85. Moulana, A. et al. Compensatory epistasis maintains ACE2 affinity in SARS-CoV-2 Omicron BA.1. Nat Commun 13, 7011 (2022).

86. Bhattacharya, T. et al. A conserved opal termination codon optimizes a temperature-dependent trade-off between protein production and processing in alphaviruses. Sci Adv 11, eads7933 (2025).

87. Tassetto, M. et al. Control of RNA viruses in mosquito cells through the acquisition of vDNA and endogenous viral elements. Elife 8, (2019).

88. Haddox, H. K., Dingens, A. S. & Bloom, J. D. Experimental Estimation of the Effects of All Amino-Acid Mutations to HIV’s Envelope Protein on Viral Replication in Cell Culture. PLoS Pathog 12, e1006114 (2016).

89. Sloan, R. D. & Wainberg, M. A. The role of unintegrated DNA in HIV infection. Retrovirology 8, 52 (2011).

90. Gerard, G. F. et al. The role of template-primer in protection of reverse transcriptase from thermal inactivation. Nucleic Acids Res 30, 3118–3129 (2002).

91. Rayaprolu, V., Ramsey, J., Wang, J. C.-Y. & Mukhopadhyay, S. Alphavirus Purification Using Low-speed Spin Centrifugation. Bio Protoc 8, e2772 (2018).

92. Loes, A. N. et al. High-throughput sequencing-based neutralization assay reveals how repeated vaccinations impact titers to recent human H1N1 influenza strains. J Virol 98, e0068924 (2024).

93. Ziegler, S. A. et al. In vivo imaging of chikungunya virus in mice and Aedes mosquitoes using a Renilla luciferase clone. Vector Borne Zoonotic Dis 11, 1471–1477 (2011).

94. Crawford, K. H. D. & Bloom, J. D. alignparse: A Python package for parsing complex features from high-throughput long-read sequencing. J Open Source Softw 4, (2019).

95. Yu, T. C. et al. A biophysical model of viral escape from polyclonal antibodies. Virus Evol 8, veac110 (2022).

96. Meng, E. C. et al. UCSF ChimeraX: Tools for structure building and analysis. Protein Sci. 32, e4792 (2023).

97. Hannon, W. W. & Bloom, J. D. dms-viz: Structure-informed visualizations for deep mutational scanning and other mutation-based datasets. J. Open Source Softw. 9, 6129 (2024).

98. Meng, E. C. et al. UCSF ChimeraX: Tools for structure building and analysis. Protein Sci. 32, e4792 (2023).

99. Chun, T. W. et al. Quantification of latent tissue reservoirs and total body viral load in HIV-1 infection. Nature 387, 183–188 (1997).

